# A thalamic hub-and-spoke circuit enables visual perception during action by coordinating visuomotor dynamics

**DOI:** 10.1101/2023.12.23.573221

**Authors:** T. Vega-Zuniga, A. Sumser, O. Symonova, P. Koppensteiner, F. H. Schmidt, M. Joesch

## Abstract

Distinguishing between sensory experiences elicited by external stimuli and an animal’s own actions is critical for accurate perception and motor control. However, the diversity of behaviors and their complex influences on the senses make this distinction challenging. Here, we uncover an action cue hub that coordinates both visual processing in the brain’s first visual relay and motor commands. We show that the ventral lateral geniculate nucleus (vLGN) acts as a corollary discharge (CD) center, integrating visual translational optic flow signals and motor copies from saccades, locomotion, and pupil dynamics. The vLGN relays these signals to correct action-specific visual distortions and refine perception, as shown for the superior colliculus and a depth estimation task. Simultaneously, brain-wide vLGN projections drive corrective actions necessary for accurate visuomotor control. Our results reveal an extended CD architecture that refines early visual transformations and coordinates actions via a distributed hub-and-spoke network enabling visual perception during action.

## INTRODUCTION

Vision is constantly challenged by the animal’s own movements ^1^. From the image alone, the origins of the sensory perturbations are ambiguous - changes in the environment (exafferent information) or consequences of the animal’s movement (reafferent information) are indistinguishable (Fig. 1A). Yet, this distinction must become operational to ensure that animals experience a behaviorally coherent perception while moving through the world ^2^. The variety and dynamic nature of animal behavior depend on complex brain-body coordination carved throughout evolution. This behavioral enrichment coevolved with an intricate battery of neural mechanisms to actively compensate for displacement of the visual world while moving. These mechanisms include the coordination of multiple senses, e.g., visual and vestibular modalities ^3^, and require an internal representation of movement commands known as efference copy or corollary discharge (CD) (Fig. 1B). The main function of CD is to filter out reafferent signals, allowing precise sensorimotor transformations needed to effectively infer the structure of the external world ^1,4^ and experience perceptual continuity ^5^. One prominent example is saccadic suppression, best understood in the primate oculomotor system, where CDs signal the direction and intensity of a forthcoming saccadic eye movement to suppress the perception of motion-induced blur ^5^. However, motion can affect visual processing in myriad ways, making effective estimation and compensation of the reafferent signal complex. This complexity is particularly evident when these corrections are distributed throughout the brain, as observed in mammals, where CDs associated with visual processing have been found primarily in thalamic and cortical regions ^1,6^.

**Fig. 1.**
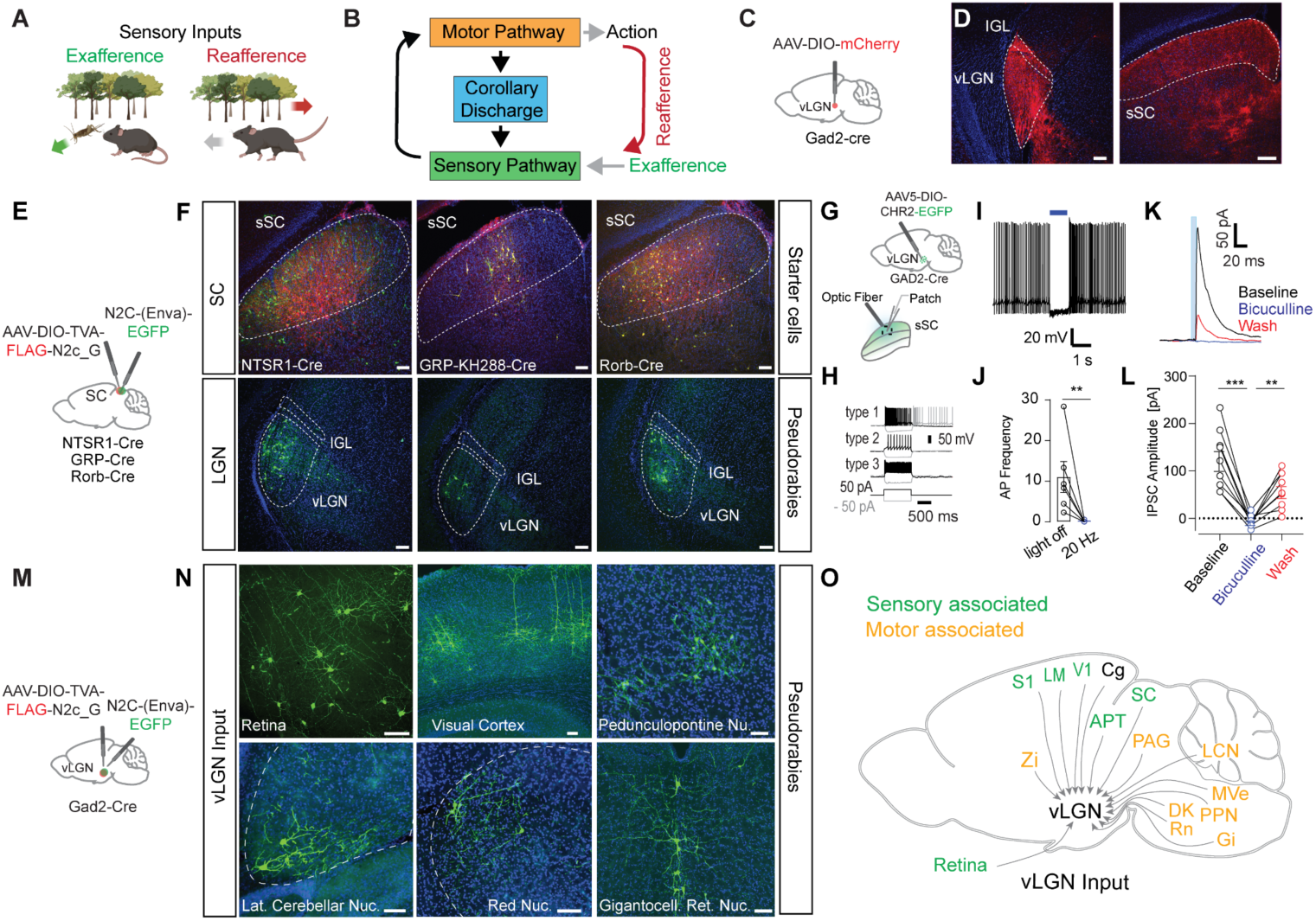
Structural and functional evidence of a thalamic corollary discharge pathway. (A) Schematic showing the difference between external visual input (exafference, green arrow) and visual input generated by the animal’s own motion (reafference, red arrow). (B) Schematic showing a sensorimotor circuit with a motor-copy signal, known as a corollary discharge (CD, blue) bridging a sensory pathway (green) and a motor pathway (orange). (C) Brain schematic showing the location of anterograde vector injection in the ventral lateral geniculate nucleus (vLGN, red dot). (D) Injection site in vLGN (left) and expression of mCherry vector in superior colliculus (right) (E) Brain schematic showing the location of retrograde transsynaptic vector injections in the sSC. (F) Coronal brain sections of the three transgenic lines. Top: sSC with starter vector expression in red. Bottom: Transsynaptic EGFP expression by the pseudorabies vectors in the vLGN. (G) Schematic brain showing a Gad2^+^ vLGN injection of ChR2 and subsequent patch clamp recordings in the sSC. (H) Intrinsic firing properties of sSC cell types in response to current steps. (I-J) Current-clamp recording(s) of spontaneous activity of sSC cell(s) suppressed by 20 Hz optogenetic stimulation (6 cells, 3 animals, paired t-test) (K) Example voltage-clamp recording of an sSC neuron in the presence of bicuculline and subsequent wash. Inhibitory currents induced by optogenetics are reversibly blocked by bicuculline (20 µM). (L) Inhibitory postsynaptic current (IPSC) amplitude quantification of all recorded neurons (9 neurons, 3 animals, Tukey’s multiple comparison test). (M) Brain schematic showing the location of retrograde transsynaptic vector injection in the vLGN. (N) Coronal brain sections of transsynaptic EGFP expression by pseudorabies vectors in sensory- and motor-associated areas. (O) Summary schematic of vLGN inputs. zona incerta (Zi), primary somatosensory cortex (S1), lateromedial area (LM), primary visual cortex (V1), cingulate cortex (Cg), anterior pretectal area (APT), superior colliculus (SC), periaqueductal gray (PAG), lateral cerebellar nucleus (LCN), medial vestibular nucleus (MVe), pedunculopontine nucleus (PPN), gigantocellular nucleus (Gi), red nucleus (Rn), darkschewitsch nucleus (DK). All scale bars: 100 µm.

The superior colliculus (SC), a prominent midbrain hub conserved across vertebrates ^7^, plays a critical role in visual sensorimotor transformations, including in humans ^8–10^. The output of the SC has been associated with CDs and has been largely studied in the context of saccadic suppression ^6^. The current view is that at least in primates, intermediate layers of the SC send ascending motor commands to the medial dorsal nucleus of the thalamus and then to the frontal eye fields in the cortex, an area known to contribute to voluntary saccades ^6^. However, the SC has been implicated in a large variety of visuomotor processes, ranging from visual perception to cognition ^11^, and some of these processes have been shown to be independent of cortical function ^12,13^. Thus, to ensure proper visual perception and visuomotor control via the SC, the SC may not only provide the motor signals required to generate the CD, but would also require processes that allow correction of a range of self-motion-induced visual distortions.

CD signals can emerge from different points along the motor pathway and affect any stage within the sensory processing stream ^1^. They can take on various functions, such as compensation, attenuation, or suppression ^14^. The last two are the best known CD signals, which are primarily mediated by inhibitory inputs. In the SC, distinct layers are targeted by long-range inhibitory projection from different brain areas ^15–18^. Among these pathways, the ventral lateral geniculate nucleus (vLGN), a retino-receptive subcortical region in the lateral thalamus ^19^, appears to be well positioned to perform CD operations. Most neurons are GABAergic ^15,16^, their connections cover all major parts of the visual brain (for details, see ^20^), and they are involved in a wide variety of tasks ranging from visual threat response behavior ^15,16^, nocifensive behavior ^21^, chromatic discrimination ^22^, optokinetic reflex ^23^ to visuomotor control ^20,24–28^. Here, we show in mice that the vLGN is a hub-and-spoke network that functions as a corollary discharge (CD) center, coordinating brain-wide visuomotor processing. The vLGN receives projections from sensory and motor-related brain regions and modulates visual responses in the earliest visual processing step, the superficial SC (sSC). It transmits CD signals from ongoing behaviors, such as locomotion, saccades, and pupil dilation, to the sSC via strong inhibitory projections. This CD signal counteracts the temporal and spatial blurring caused by animal behavior and is thus a fundamental component of proper visual perception. Accordingly, animals with a disrupted vLGN show deficits in a task that requires the integration of vision and action. Beyond the sSC, we show that the vLGN also projects to several motor control related areas, forming a distributed processing spoke-like system that coordinates early sensory processing and motor control. Optogenetic activation of the vLGN produces stereotyped corrective movements, whereas targeted suppression disrupts the precision of these behaviors. Taken together, our data show that the vLGN plays a critical role in coordinating visuomotor transformations during action, resembling a distributed feedback control system.

## RESULTS

### The vLGN forms a hub integrating disperse sensory and motor areas

The vLGN/IGL complex is a prethalamic nucleus composed of mainly inhibitory neurons ^15,16^. Recent work has shown that the vLGN projects across collicular layers (Fig. 1C,D), modulating collicular processing by relaying information of threat levels or feature related visual signals ^16,29^. However, the relative contribution of the vLGN and IGL, in particular to the visual recipient layers, remained contested and the specificity of these projections unknown. To determine the relative contributions of vLGN and IGL^16^ to sSC cell types, we used highly neurotropic N2c rabies virus vectors ^30^. We retrogradely labeled the presynaptic neurons of predominantly glutamatergic sSC neurons using the Ntsr1-GN209 (wide-field), Grp (narrow-field), and Rorb-Cre (stellate and other neurons) lines (Fig. 1E,F)^31^. In all cases, we observed predominantly vLGN labeling, with few or no labeled cells in the IGL, and no labeled neurons in the dLGN (Fig. 1F). To confirm that the vLGN exerts direct inhibitory control of sSC, we optogenetically stimulated Gad2^+^ vLGN terminals expressing channelrhodopsin-2 (ChR2) in combination with *in vitro* whole-cell recordings of sSC neurons (Fig. 1G-L). First, we verified that we sampled across a range of cell types, by characterizing the physiological properties of each cell in response to current injections ^31^ (Fig. 1H). In neurons that fired spontaneously, sustained optogenetic activation of vLGN terminals led to a near abolishment of spikes (Fig. 1I,J). In voltage clamp, short optogenetic stimulation (5 ms) reliably led to large, GABA_A_-dependent, inhibitory postsynaptic currents (124 ± 21 pA) (Fig. 1K,L). Overall these data show that the vLGN is well-positioned to exert a strong modulatory influence on the first visual relay, and likely also across the wide-spread sensory and motor projecting areas ^32^.

Next, using N2c rabies vectors, we mapped the areas providing direct input to Gad2^+^ vLGN neurons (Fig. 1M). In addition to previously characterized inputs from the retina ^33^, we observed direct projections from a wide-spread network of subcortical and cortical areas. We found inputs from cortical areas such as visual and cingulate cortical areas, involved in emotional and cognitive control ^34^, as well as fear and stress responses ^35^ (Fig. 1N,O; Fig. S1). We also observed inputs from several pre-motor and motor nuclei, such as red nucleus, pedunculopontine nucleus, gigantocellular reticular nucleus, and lateral cerebellar nucleus, which are known to be required for motor coordination (Fig. 1N, O, Fig. S1). Interestingly, some of these areas project indirectly or directly ^36,37^ to the spinal cord, indicating that copies of motor commands are transmitted directly to the vLGN.

These results demonstrate that the vLGN is situated in the center of an extended network linking motor and sensory areas with inhibitory connections, leading to the hypothesis that it provides CD-type control of visual centers during behavior. Notably, its projection targets prominently include the ipsilateral sSC as a sensory example.

### The vLGN modulates visual response properties in the early visual system

To explore the modulatory influence of vLGN on visual processing, we simultaneously recorded visually-evoked responses across different layers of the SC, using silicon probes in head-fixed awake-behaving mice, and optogenetically activated the vLGN (Fig. 2A, B). For this purpose, we used the same viral delivery approach and Gad2-cre mice as for *in vitro* physiology (Fig. 1G) and subsequently implanted an optical **λ**-fiber for optogenetic stimulation (Fig. 2C, D). First, we determined the recording depth using visually evoked responses and current source density (CSD) analysis (Fig. 2E, left), which was aligned with the histological reconstruction of the probe position (Fig. 2D). When visual stimuli were replaced by optogenetic pulses in the vLGN, we observed an inversion of the current source, reflecting the inhibitory nature of the thalamic projection (Fig. 2E, right). The response depth of the visual and optogenetic responses overlapped, but the optogenetic CSDs reached further into the intermediate layers (Fig. 2F) as expected from the anatomical projection. Visual responses peaked at ∼ 60 ms after the visual stimulus onset likely due to the speed of the phototransduction, whereas optogenetic responses had a peak-latency of a few milliseconds (Fig. 2G). To quantify how these pronounced inhibitory dynamics influence visual responses in the sSC, we combined small visual flashes (10 ° of visual angle, duration 200 ms or 1 s) centered in the receptive fields of the recorded sSC units with interspersed and randomized vLGN optogenetic stimulation (Fig. 2H). Next, we determined the units that were visually and optogenetically responsive, independent of variations in retinotopic position, using parameter-free stimulus-evoked responsiveness tests to directly examine interactions. 66 % (376 out of 571) of all recorded units were visually responsive, out of which 67 % (253 out of 376) were responsive to optogenetic stimulation. On average, this population showed a mild modulation of baseline firing (Fig. 2H, middle), but a strong and effective suppression of visually evoked responses (Fig. 2H, right, I), reducing the maximum firing rate by ∼60% (Fig. 2H, bottom, I). Since the relative timing of the first spike has been shown to be an effective retinal code ^38^, we next asked whether feedforward suppression would affect spike timing. We tested spike timing precision on the subset of units that did not have complete suppression and compared the latency of the first spike to a visual stimulus between control and optogenetically stimulated trials. On average, optogenetic stimulation had a small effect, delaying the first spike by 3.9 ms (Fig. 2J), indicating that vLGN inhibition in the SC mostly affects the rate but not the timing of the first spike. To determine the kinetics of inhibition, we sorted the trials by the relative onset time of the optogenetic stimulation (Fig. 2K). vLGN suppression was largely transient, was strongest when visual and optogenetic stimuli overlapped, and lasted for approximately 100 ms after the offset of the optogenetic stimulus (Fig. 2L). Consistent with the extension of vLGN projection to lower SC layers ^15,16^, we observed similar suppression in intermediate SC layers (Fig. S2). Finally, we tested whether vLGN activation would modulate sensory properties such as the spatiotemporal receptive fields (RF) of sSC neurons. We mapped the 1-dimensional RF using vertical bars appearing at random horizontal locations interleaved with optogenetic stimulation. As shown previously ^29^, we observed a sharpening of the spatial RF (Fig. 2M,N).

**Fig. 2.**
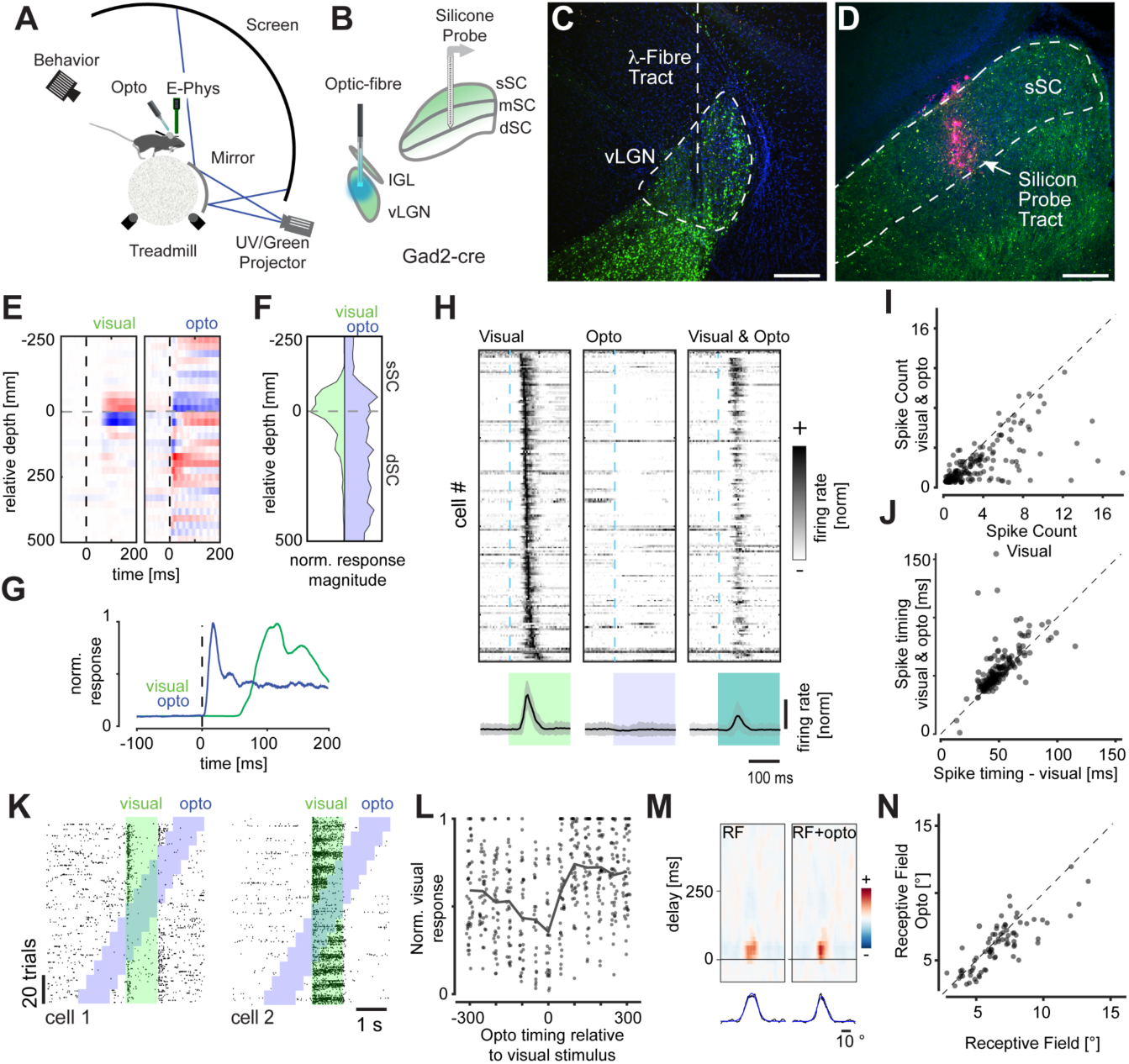
The vLGN modulates the rate, not the timing of early visual processing *in vivo*. (A) Schematic of the electrophysiology setup, with mice head restrained while running on a spherical treadmill with visual stimulation on a panoramic dome. (B) Schematic of ChR2 virus infection in Gad2^+^neurons in the vLGN and placement of the recording electrode. (C) Confocal image of the vLGN and the optical fiber track for optogenetics. (D) Confocal image of the SC with the DiI-labeled track of the recording electrode. (E) Current source density (CSD) analysis for an example recording for visual flash stimuli (left) and optogenetic stimulation (right). Black vertical dashed line indicates stimulus onset. (F) Average contours of normalized CSD over depth (26 recordings, 5 animals). (G) Temporal profile of CSD activation for the recording in (E). (H) Sorted and normalized firing responses of sSC units to visual, optogenetic, and combined stimulation (top) with their respective population mean responses (bottom shows mean (black) ± SD (shaded grey); n = 301, 26 recordings, 5 animals). (I) Quantification of optogenetic activity suppression, spike count within 0.2 s after the flash onset (n = 301, 26 recordings, 5 animals. Two-sided Wilcoxon signed rank test, p < 0.001, mean ± SD of spike count difference −1.12 ± 2.15). (J) Analysis of the optogenetic influence on visually evoked spike timing (n = 301, 26 recordings, 5 animals. Two-sided Wilcoxon signed rank test, p < 0.001, mean ± SD of spike timing difference 3.9 **±** 11.9 ms) (K) Spike raster plots of visually and optogenetically responsive sample units of the superficial SC (sSC). Visual stimulus epochs (green) are interleaved with optogenetic stimulation (blue). (L) Quantification of the duration of inhibition kinetics, n = 81, 7 recordings, 4 animals. (M) Horizontal receptive field of an example neuron during random vertical bar stimulus (left), with optogenetic stimulation (right) and their horizontal profiles (bottom, data (black) and gaussian fit (blue)). (N) Quantification of the difference of sizes of the centers of the receptive fields with and without optogenetic stimulation. (n = 79, 10 recordings, 3 animals. Two-sided Wilcoxon signed rank test, p < 0.001, mean ± SD of size difference of the center of receptive fields −0.53 ± 1.23 degree). Scale bars in C & D: 100 µm.

Collectively, these data show that the vLGN acts as a feedforward inhibitory hub capable of strongly reducing visual responses across the sSC, while leaving onset of timing largely unaffected, thus effectively sharpening visual responses in both spatial and temporal domains.

### vLGN is a central hub for corollary discharges

To investigate the scenarios in which the vLGN modulates sSC visual processing, we analyzed the response properties of vLGN axon terminals in the sSC (Fig. 3A). We introduced the calcium indicator axon-GCaMP6s ^39^ into Gad2^+^ cells by infecting the vLGN (Fig. 3B-D, n = 4 mice), resulting in homogeneous axonal GCaMP expression throughout the SC (Fig. 3C). Subsequent implantation of a cranial window over the SC allowed observation of activity in axonal terminals from the thalamus using two-photon calcium imaging in awake, behaving mice. In addition, retinal terminals in the SC were recorded to qualitatively compare their response characteristics (Fig. S3). We used a variety of visual stimuli to assess bouton response properties, with interleaved periods of uniform light level (gray dome screen) to assess the activity associated with spontaneous animal behavior.

**Fig. 3.**
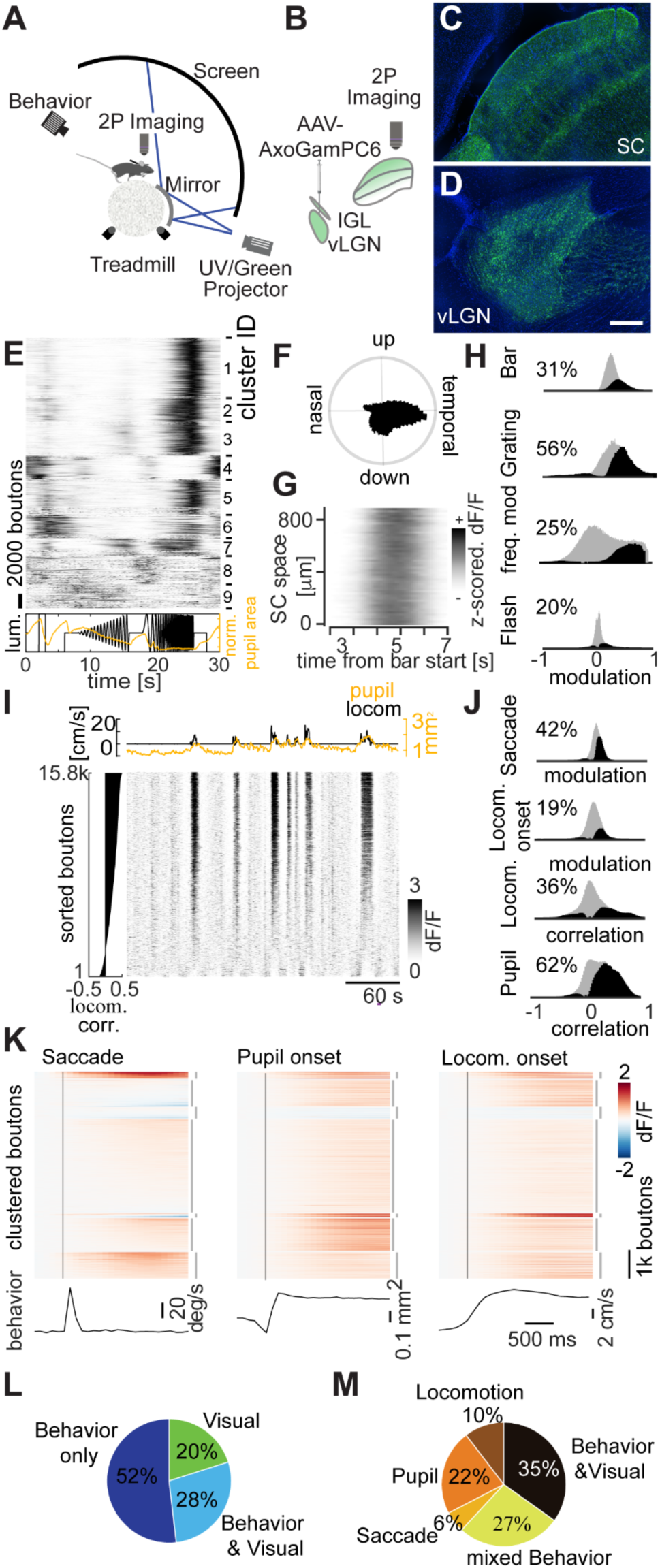
vLGN projections to the SC are strongly modulated by behavior. (A) Schematic of the multi-photon imaging setup with head-restrained mice running on a spherical treadmill and panoramic dome visual stimulation. (B) Schematic of axon-GCaMP6s viral expression in vLGN Gad2^+^ neurons. (C) Confocal image of the axonal terminals in the sSC and (D) the infection area in the vLGN, sagittal view. Scale bar: 100 µm. (D) Cluster-sorted, averaged and z-scored responses (top) to full field luminance chirps (bottom, black line) of vLGN boutons in the sSC and average modulation of normalized pupil area (bottom, orange line). (E) Polar histogram of direction-selective (p < 0.01, shuffle test, n = 31620 (26%)) boutons’ preferred direction in response to full field drifting gratings. (F) z-scored and spatially binned (20 µm) vLGN bouton responses across SC space to a moving bar moving in nasal-temporal direction. (G) Histograms of modulation indices of vLGN bouton population (gray) and significantly modulated boutons (black; two-sided Wilcoxon signed rank test, p < 0.01) to visual stimuli. (H) Raster plot of example recording of vLGN boutons sorted by correlation with locomotion (left) and respective locomotion (black) and pupil area (orange, top) traces (I) Histograms of modulation indices (saccades, locomotion onset) or correlation coefficients (locomotion speed, pupil area) of vLGN/IGL bouton population (gray) and significantly modulated boutons (black & percentages) to behavioral parameters. p < 0.01, two-sided Wilcoxon signed rank test, shuffle test for correlations (see methods). (J) Average activity modulation of vLGN boutons (n = 131871, 17 recordings, 5 animals, only including recordings with at least 10 trials per each condition) aligned to saccades (bottom: average eye movement speed), rapid pupil expansion onset (bottom: average pupil area) and locomotion onset (bottom: average forward locomotion speed). Time of behavioral event indicated by vertical black line. Boutons sorted by k-means clustering (interrupted vertical lines on the right, k = 7) joint activity profiles. (K) Pie chart of boutons significantly modulated by any of the visual stimuli tested (H) and/or significantly modulated by/correlated to any of the behavioral parameters (J); p < 0.01, Bonferroni corrected for multiple comparisons. (L) Pie chart of boutons significantly modulated by/correlated to any of the behavioral parameters (J) individually, removing purely visually responsive neurons (n = 176282, 38 recordings, 6 animals); p < 0.01, Bonferroni corrected for multiple comparisons. n = 220021, 38 recordings, 6 animals, if not stated otherwise.

First, we tested bouton responses to a “chirp” stimulus ^40^, which involves full-field modulations of light intensity (Fig. 3E). Among all vLGN boutons meeting the inclusion criteria (see methods), some showed robust responses to full-field flashes, as previously reported ^41^. However, the majority showed their most pronounced responses to rapid frequency modulations, which can occur during locomotion, such as running underneath nearby vegetation. Retinal boutons, on the other hand, showed a wider range of feature selectivities to the stimulus (Fig. S3E). Next, we determined directional selectivity to full-field moving grating stimuli, which in vLGN boutons showed a striking preference for temporal grating motion (Fig. 3F), as would be seen during forward locomotion. In contrast to retinal ganglion cell (RGC) responses ^42^, the use of a random checkerboard or sparse dot stimulus did not yield robust receptive field estimates for vLGN boutons (not shown). Nevertheless, we wanted to determine the retinotopic organization of the terminals in the sSC. To this end, we tested whether a visual bar moving slowly (22.5 °/sec) across the visual field elicited sequential activations that were anatomically localized. Retinal synapses showed a distinct spatial representation (Fig. S3G), whereas vLGN terminals showed global responses when a bar crossed visual space, consistent with electrophysiological data indicating large spatial receptive fields ^41^ (Fig. 3G). Overall, approximately half (48 %) of the included vLGN boutons (RGC: 93 %) responded significantly (p < 0.01, Bonferroni corrected) to at least one of the visual stimuli presented (Fig. 3H).

During gray screen periods, activity in retinal boutons was weak and sparse (Fig. S3I), whereas calcium signals in vLGN boutons varied strongly (Fig. 3I). Compared to the retinal terminals, vLGN boutons activity had a high degree of synchrony (on average 18% variance explained by population mean for vLGN, 3% for RGCs; Fig. S4A, B), indicating that the vLGN is transmitting a global signal. We found that activity peaks often coincided with locomotor bursts and thus compared vLGN bouton signals with behavioral parameters. Indeed, many vLGN boutons significantly changed their activity at saccade, pupil dilation and locomotion onset (Fig. 3J, K). Confirming previous results ^43^, retinal inputs to the sSC were also modulated by behavior during gray periods, with a quarter of retinal boutons significantly affected by at least one of the measured behavioral parameters (Fig. S3J-L, S4C). However, vLGN bouton activity was coupled to behavior to a much greater extent and proportion, with 80 % of boutons significantly and mostly positively modulated by behavior (Fig. 3L), in stark contrast to retinal axons (Fig. S3L). Interestingly, ∼38 % of behaviorally modulated vLGN boutons were specifically modulated by only one behavioral parameter (Fig. 3M), suggesting that the vLGN is composed of specific cell-types carrying distinct information, in line with single-cell sequencing data of the vLGN reporting a large neuronal diversity ^44^. Since the dynamics of locomotion and pupil size can be coupled (median correlation across recordings c = 0.24, n = 67 recordings, Fig. S4F), we next tested whether vLGN bouton activity coupling to each behavior is independent. We first tested the relative neuron-behavior cross-correlation timing, determining that correlation to running speed was maximal when vLGN bouton activity was shifted to slightly preceding the behavior (Fig. S4D), in contrast to pupil area, where minimally delayed neuronal activity led to maximal correlations (Fig. S4E). To further uncouple pupil size and locomotion in our analyses, we compared the difference of correlation of vLGN and retinal activity with pupil size overall and during stationary epochs only. We found that in stationary periods pupil size correlations were significantly lower for both bouton populations, but to a far larger extent in vLGN boutons (mean difference ± SD = 0.084 ± 0.138) than RGC boutons (mean difference ± SD = 0.005 ± 0.093), indicating that locomotion and pupil dynamics exert an independent influence on vLGN boutons (Fig. S4G-I). In conclusion, the vLGN to sSC projections are activated by cross-modal signals, i.e. visual and behavioral. This activity is well positioned to provide a potent inhibition of visual signals when changes in the retinal image are expected to occur due to self-motion, e.g. to counteract luminance changes during pupil dilation or motion blur during locomotion or saccades.

### The vLGN coordinates visual and motor signals

Our anatomical and physiological data (Fig. 1-3) suggest that the vLGN acts as a feedback controller, anticipating the effects of movement on the animal’s visual input. Thus, this pathway should be required to correct for movement related visual blur, e.g., being involved in saccadic suppression, and for maintaining perceptual stability during behavior. We tested this hypothesis by comparing vLGN responses to saccades while displaying a stationary structured background with visually evoked responses to “pseudosaccades” (movements of the displayed pattern that would mimic the visual input perceived during saccades, Fig. 4A) ^45^. As expected, retinal boutons in the SC responded similarly, irrespective if the eye or visual stimulus moved (Fig. 4B). In contrast, vLGN boutons were mainly sensitive to real saccades (Fig. 4C), showing a clear preference for voluntary eye movement saccades over pseudosaccades (Fig. 4D). Next, to directly test whether the vLGN/sSC network functions as a feedback control loop, we chronically blocked bilateral Gad2^+^ vLGN output by expressing tetanus toxin light chain (TeLC) ^46^ (Fig. 4E, F) and recorded sSC electrophysiologically as previously (Fig. 1, 2). First, we confirmed *in vitro* that TeLC blocks synaptic release (Fig. S5A-C). We then verified *in vivo* that behaviorally independent visual responses in SC were largely unaltered (Fig. S5D-F). Next, we tested if the vLGN is required to reduce the motion blur relayed by the retina during saccades while viewing a high-contrast screen. Indeed, in animals in which the vLGN output was blocked, sSC responses to saccades were on average 100 ms longer (Fig. 4G-H) and showed increased average firing rates in the first 200 ms (Fig. 4I). Thus, such action-induced feedback can be thought of as a neural mechanism that reduces the effective visual exposure time during actions.

**Fig. 4.**
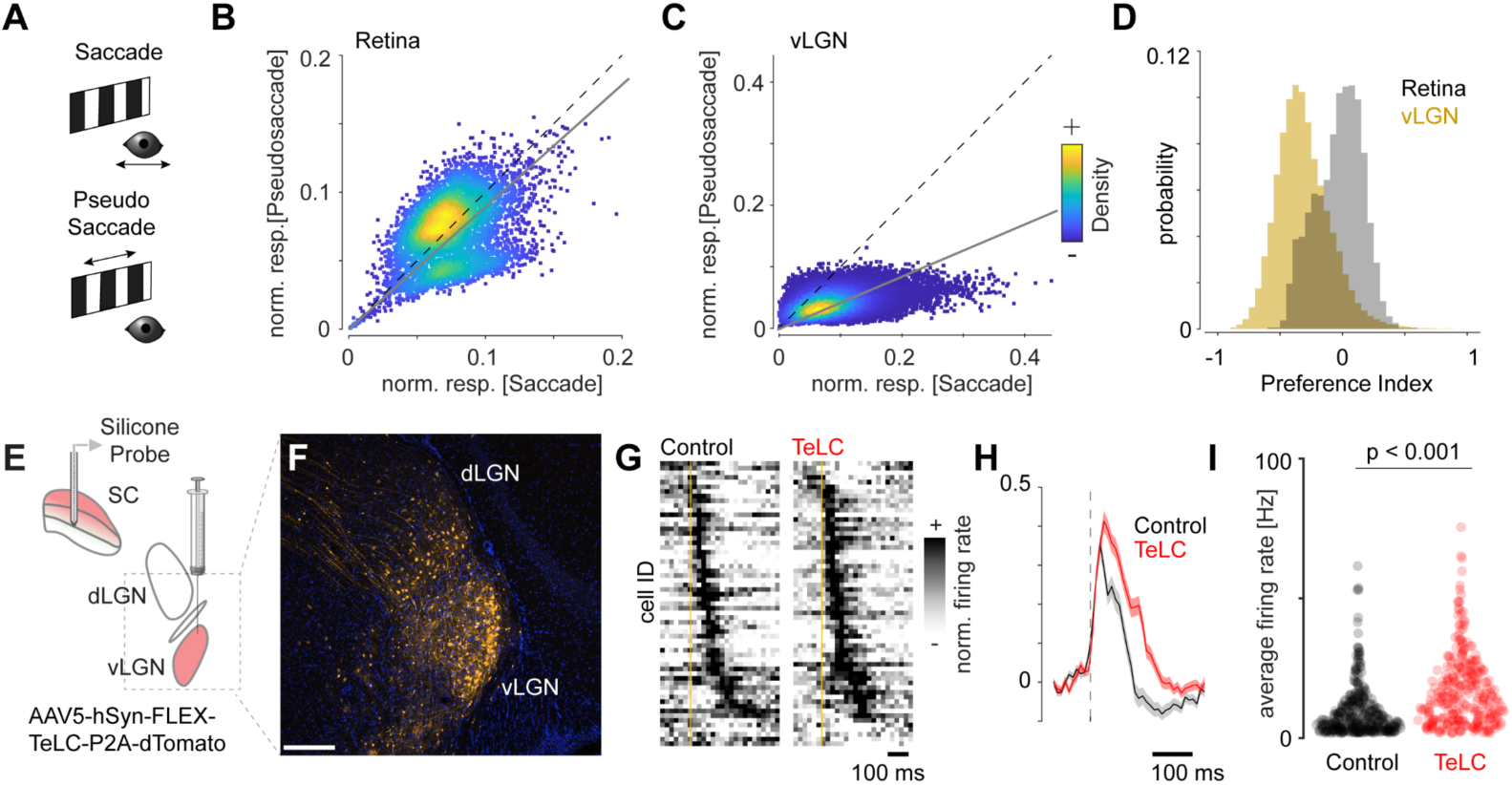
The vLGN suppresses behaviorally induced sensory blur. (A) Spontaneous saccades on a stationary grating or checkerboard (“saccades”, top, black) and saccade-like shifts of the displayed texture during stationary eye periods (“pseudosaccades”, bottom, green) induce comparable image shifts. (B) Density-colored scatter plot of normalized responses of individual retinal boutons comparing pseudosaccades and saccades, showing stronger responses to pseudosaccades (linear fit: gray line, slope = 0.89, p < 0.001, paired, two-tailed Wilcoxon signed-rank test, n = 5174). (C) Same as B but for vLGN boutons with a strong preference for saccadic motion (linear fit: gray line, slope = 0.42, p < 0.001, paired, two-tailed Wilcoxon signed-rank test, n = 51996). (D) Histograms of the bouton-wise pseudosaccade preference index for RGC (gray) and vLGN (yellow) bouton responses as in (B, C) with vLGN bouton preference shifted to the negative compared to RGC (p < 0. 001, two-tailed Wilcoxon rank sum test, vLGN: mean ± SD = −0.310 ± 0.214, RGC: mean ± SD = −0.030 ± 0.187). (E) Schematic of virally mediated TeLC expression in vLGN and SC extracellular recording. (F) Confocal micrograph of TeLC-tdTomato expression (orange) in thalamus of Gad2-cre mice. Scale bar: 100 µm. (G) Normalized peristimulus time histogram (PSTH) of mean saccade-triggered responses in control (left) and TeLC (right) mice (control n = 370, 8 recordings, 5 animals; TeLC n = 295, 6 recordings, 4 animals). (H) Normalized population-averaged saccade responses and standard error of the mean (SEM) (shading). (I) Average firing rate for the first 200 ms after saccade onset (p < 0. 0001, KS-Test)

### vLGN feedback is required for visual perception in action

Our previous results suggest that the vLGN is required for the brain’s ability to interpret visual signals during motion. Recently, it has been shown that mice can use monocular and binocular cues to estimate depth ^47^ and that monocular perception requires motion for proper depth estimation ^48^. We therefore decided to test depth perception using a classic visual cliff paradigm (Fig. 5B), which relies on monocular visual activity from the lower visual field, where there is minimal binocular overlap ^47,48^. For these experiments, to minimize tactile sensory input, we clipped the whiskers of TeLC-mediated (Fig. 5A) bilaterally vLGN-blocked and control mice, placed the animals individually on the platform, and recorded their behavior. Control and TeLC mice showed no difference in their average running speed while exploring the arena (Fig. 5C), indicating that chronically blocking vLGN did not grossly change locomotor and general exploratory behavior. Control mice showed a strong preference for the platform, but also roamed around the edges of the arena, possibly touching the walls with their bodies (Fig. 5D, top). We therefore restricted our analyses to the central part of the arena. vLGN-blocked mice showed a strong reduction in cliff avoidance (Fig. 5D, E). In line with these findings, we observed that control animals more frequently aborted movements out of the platform, compared to vLGN-blocked mice (Fig. 5D, G), suggesting that vLGN-blocked animals have difficulty judging depth. This impairment is linked to an increase in visual blur during motion caused by various behaviors when the vLGN is blocked (Fig. 4), affecting motion parallax computation. Finally, TeLC expression in adjacent medial thalamic areas, including the zona incerta, did not affect cliff avoidance behavior, indicating that the vLGN is specifically required (Fig. S5G-J). Taken together, these results show that visuomotor processing and perception during self-generated motion are impaired in the absence of proper vLGN function.

**Fig. 5.**
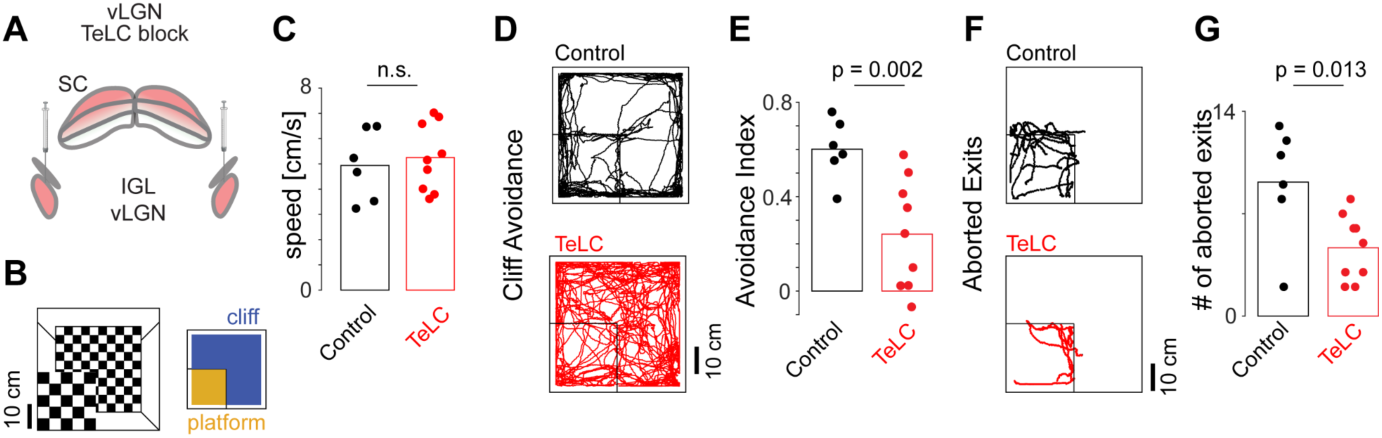
The vLGN modulates visual percepts during voluntary movements. (A) Schematic of the bilateral TeLC infection of vLGN. (B) Schematic of the visual cliff setup (left) and areas used for subsequent quantification (right). (C) Average running speed of control and TeLC mice in the arena (6 control and 9 TeLC animals). (D) Examples of running trajectories during the first 10 min for control and TeLC mice. (E) Cliff avoidance quantification (see methods) of (D) per animal (Wilcoxon rank sum test). (F), as (D), but only showing trajectories for aborted exits (2 s before and after a detected crossing with the subsequent reversal of the motion velocity, see methods) and (G) the corresponding quantification (Wilcoxon rank sum test).

### Distributed connections from the vLGN drive corrective actions

Several classical studies indicate a wide-spread connectivity architecture, yet have not distinguished vLGN projections from those in the intergeniculate leaflet (IGL) ^32^ and zona incerta (ZI). To determine the brain-wide projections that arise specifically from the vLGN we performed anterograde labeling experiments (Fig. 6A). We isolated vLGN neurons by transsynaptically labeling visual cortex (VC) target neurons, which in the thalamus includes the vLGN and the dorsal LGN (dLGN), but excludes IGL (Fig. 6B). We did not observe any retrogradely labeled L5 pyramidal cells in the VC (Fig. S7A,C). Corroborating our previous findings (Fig. 1), vLGN projections to the sSC were directed to the lower layers of the sSC (Fig. 6C middle/top), where most somata of retinorecipient neurons are located ^31^. We also observed projections to the intermediate layers of the SC ^15,16^, but most prominently a wide projection pattern to a range of subcortical sensory and motor related areas. Targeted sensory areas are located in the thalamus (lateral posterior nucleus (LP), nucleus reuniens, contralateral dLGN and vLGN, lateral habenula) and midbrain (posterior pretectal nucleus, anterior pretectal nucleus, olivary pretectal nucleus, periaqueductal gray) (Fig. 6C, Fig. S6). Our results indicate that the required action induced sensory signals may originate partially from the vLGN. Projections to motor related areas include midbrain and hindbrain structures (Fig. 6C, S6). For example, the pretectal olivary nucleus, an area known to directly activate the Edinger-Westphal nucleus, in turn the strongest regulator of pupillary constriction ^49^. It also projects to the pons and inferior olive, involved in visuo-motor coordination ^50^, deep mesencephalic nucleus, known as an output center of basal ganglia ^51^, and to the red nucleus, an area suggested to be involved in fine motor coordination ^52^. These projection patterns can be recapitulated by targeting Gad2^+^ cells in the vLGN/IGL (Fig. S7D,E) ^32^, indicating that GABAergic neurons in vLGN are the major contributors to these projections.

**Fig. 6.**
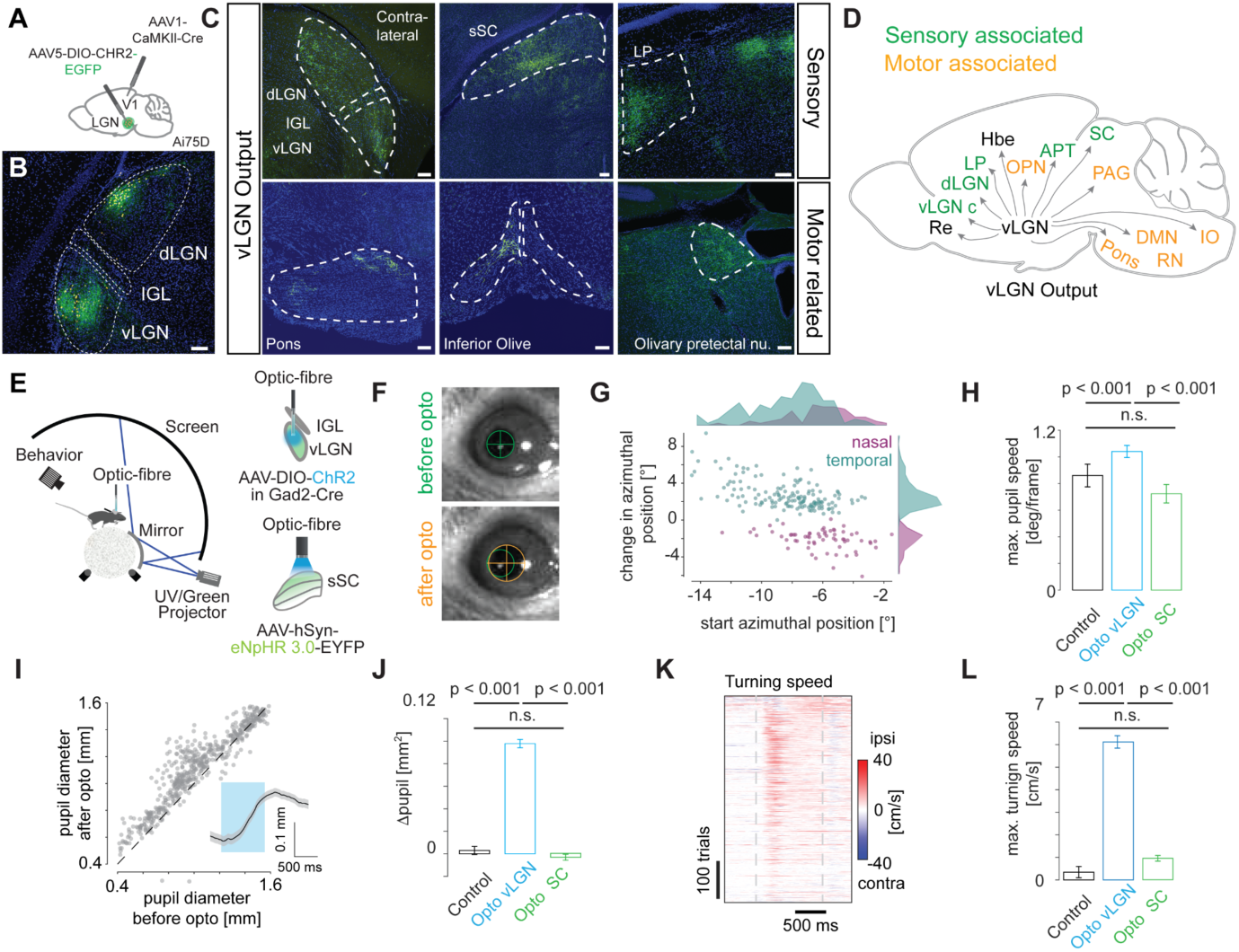
Brain-wide vLGN projections fine-tune behavior. (A) Brain schematic showing the location of anterograde transsynaptic vector injection. Primary visual cortex (V1, red dot) and ventral lateral geniculate nucleus (vLGN, green dot) in Ai75D mice, which express nuclear-localized tdTomato following Cre recombinase exposure. (B) Brain section showing channelrhodopsin (ChR2)-EGFP expression after Cre-dependent vector injection in the vLGN. (C) Expression of ChR2 in vLGN terminals in sensory and motor associated areas. (D) Schematic summary of vLGN outputs. Nucleus reuniens (Re), ventral lateral geniculate nucleus contralateral (vLGN c), dorsal lateral geniculate nucleus (dLGN), lateral posterior nucleus (LP), habenula (Hbe), olivary pretectal nucleus (OPN), anterior pretectal area (APT), superior colliculus (SC), periaqueductal gray (PAG), inferior olive (IO), deep mesencephalic nucleus (DMN), red nucleus (Rn). (E) Left, schematic of the setup with head-restrained mice while running on a spherical treadmill. Right, schematic of vLGN (top) and sSC (bottom) optogenetic stimulation. (F) Example images of an optogenetically induced saccade. Top, before optogenetic stimulation, bottom, after. Green and orange circles demarcate the size and position of the pupil, before and after stimulation, respectively. (G) Magnitude and direction of optogenetically induced saccades (only saccades that exceeded a speed threshold were plotted - see methods, 211 repetitions, 3 animals, 9 recordings). (H) Quantification of optogenetically induced pupil velocity (KS test, control: 62 repetitions, 3 recordings, 2 animals; opto vLGN: 211 repetitions, 9 recordings, 3 animals, opto SC: 185 repetitions, 3 recordings, 3 animals). (I) Optogenetically induced pupillary dilation. Inset: average kinetics (p<0.001 Wilcoxon signed rank test, 544 repetitions, 11 recordings, 4 animals). (J) Quantification of optogenetically induced pupil change (same as L, KS test, control: 229 repetitions, 3 recordings, 2 animals; opto vLGN: 546 repetitions, 11 recordings, 4 animals, opto SC: 779 repetitions, 3 recordings, 3 animals). (K) Optogenetically induced turning speed. Gray dashed lines represent the start and end of the optogenetic stimulation (546 repetitions, 11 recordings, 4 animals). (L) Quantification of optogenetically induced turning speed for control, vLGN and SC.

Our results show that the vLGN acts as a feedback controller, fine-tuning sensory signals corrupted by motion blur (Figs. 1-5). However, anatomical projections to several motor-related areas suggest that the vLGN functions on a more global scale, coordinating visual and motor processes simultaneously. While motion blur corrections are relevant for any action and should be corrected in the periphery, other corrections may require specificity, especially if the actions occur on different time scales (e.g. saccades are ballistic movements, pupil dilation is continuous). Thus, we reasoned that optogenetic activation of vLGN pathways might reveal motor corrections related to the behaviors associated with its neuronal activity (Fig. 3). To test this, we analyzed the behavioral responses to vLGN activation in the previously recorded animals (Fig. 2), focusing on locomotion, saccadic eye movements, and pupil size modulation (Fig. 6E-L). Upon optogenetic activation, mice robustly changed their gaze (Fig. 6F), which had a small corrective movement of ∼3 ° on average and tended to increase in magnitude when the eye was further away from the central position before stimulation (Fig. 6F-H). We also observed a robust increase in pupil size upon optogenetic stimulation, independent of the initial dilation (Fig. 6F, I-J). Similarly, their walking direction and rotation of the spherical treadmill were strongly toward the side of optogenetic stimulation (Fig. 6K, L, Fig. S8A,B). These behaviors do not appear to be mediated by the sSC, as optogenetic inhibition using panneuronal eNpHR3.0 of large parts of the sSC in control experiments showed no behavioral effects (Fig. 6E-L, Fig. S8). These results show that beyond modulating visual signals, vLGNs globally influence sensorimotor transformations, coordinating sensation and action.

### vLGN is required for the control of corrective actions and arousal

Optogenetic vLGN activation can drive corrective movements consistent with its functional repertoire (Figs. 3 & 6), suggesting that the vLGN is part of a finely tuned feedback control system that adapts to the behavioral needs of the moment. To test the role of the vLGN as a global feedback controller that coordinates visual and behavioral transformations in parallel, we quantified fine-grained motor deficits in animals in which the vLGN was blocked by targeted bilateral TeLC expression (Fig. 7A). First, we tested if vLGN-blocked animals would have deficits in visuomotor transformations, such as with the optokinetic reflex. For this, we used a sinusoidally moving random checker and measured tracking accuracy by eye movements ^53^ (Fig. 7B). On average, control and vLGN-blocked animals could track the sinusoidal movement to a similar extent (Fig. 7C). However, while control animals showed a large number of saccades in the direction of stimulus motion, on average 0.4 saccades per trial, vLGN-blocked animals had essentially no saccades during non-running epochs (Fig. 7D,E). The functional interpretation of such corrective actions remains elusive, but they appear to reveal mechanisms required to improve the accuracy and stability of eye movements similar to those described in humans ^54^. Next, we tested whether the sensory evoked pupil constriction was affected by measuring the pupil constriction reflex in animals with TeLC-mediated vLGN block, compared to control animals (Fig. 7F). We observed a drastic decrease in the constriction rate in vLGN blocked animals (Fig. 7G, H). Next, we tested whether pupil dynamics were altered in relation to locomotion state. Both during stationary (Fig. 7I) and running periods (Fig. 7J, K), pupil sizes were significantly smaller in vLGN-blocked animals, never reaching a fully dilated state. These results emphasize that the fine control of visually driven and running/arousal coupled oculomotor behavior requires signals relayed by the vLGN, and suggest that the vLGN synchronizes both visual processing and animal behavior in real time (Fig 8B).

**Fig. 7.**
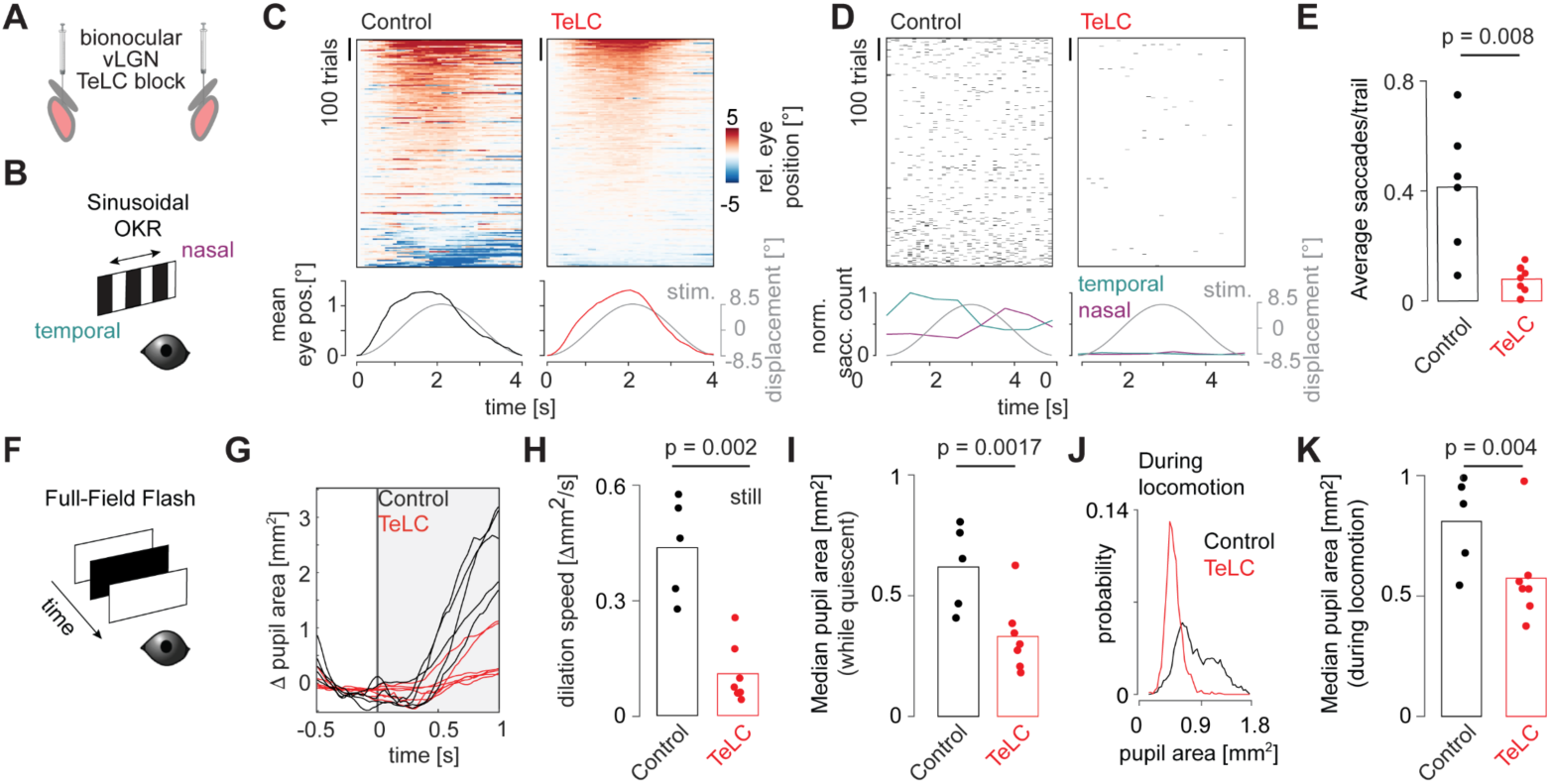
The vLGN activity is necessary for the coordination of behaviors and arousal. (A) Schematic of bilateral viral TelC expression in Gad2^+^ neurons in the vLGN/IGL complex. (B) Schematic of optokinetic reflex paradigm. (C) Top. Sorted eye positions for control and TeLC animals in trials where animals were not locomoting. Each trial is zeroed to the starting position. Bottom. Average eye position plotted next to the stimulus displacement. Note the absence of saccadic movements in TeLC animals compared to controls. (D) Top. Saccadic events (displacements > 2.5 deg/frame), trial sorted as in (C) Bottom. Normalized saccadic counts for temporal and nasal saccades. Note: Control animals correct the smooth optokinetic tracking with saccadic events, a behavior absent in TeLC animals. (E) Average saccadic events per trial per session (Wilcoxon rank sum test, control: 5 animals, 6 sessions, TeLC, 6 animals, 7 sessions, same for C and D) (F) Schematic of pupillary reflex paradigm. (G) Average stimulus evoked pupil dilation per animal (control 5 animals, TeLC 7 animals). (H) Stimulus evoked dilation speed from L (Wilcoxon rank sum test, same for I and K) (I) Median pupil area during still periods for control and TeLC animals. (J) Pupil area distribution during running for control and TeLC animals. (K) Median pupil area during running periods for control and TeLC animals.

**Fig. 8.**
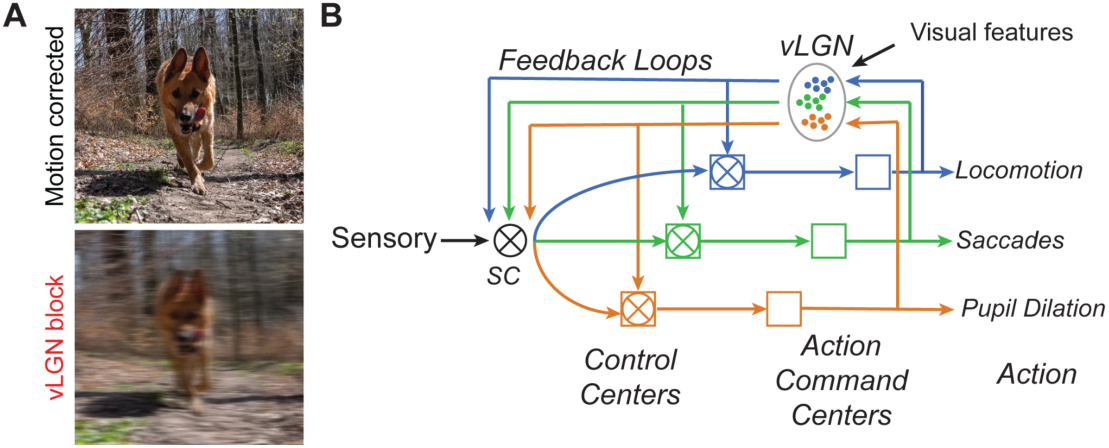
Summary of the vLGN function in coordinating sensation and action. (A) Illustrative motion blur correction. (B) Schematic of the vLGN hub-and-spoke feedback loops that coordinate vision and behavior.

## DISCUSSION

All animals require mechanisms to distinguish sensory inputs caused by their own movements (reafferent) from sensory inputs caused by sources in the outside world (exafferent) (Fig. 1A). In the visual system, several mechanisms have been proposed to support such perceptual tasks. These start at the retina ^55,56^, acting in concert with ocular stabilization mechanisms such as the vestibulo-ocular reflex (VOR) ^57–59^. However, despite this precisely tuned motor feedback system, there are several types of self-induced visual motion that cannot be stabilized, e.g., visual motion during translational movements. An additional strategy for providing corrective signals is via corollary discharges (CD) ^1^, neural mechanisms that involve the transmission of a copy of motor commands or signals from a brain region involved in generating motor output to brain regions processing sensory information. Thus, a CD can provide a predictive signal about the expected sensory feedback resulting from the motor command. In the visual system, CD has been extensively studied in the context of saccades, rapid eye movements that would temporarily blur visual perception. To minimize this perceptual blurring, a process called saccadic suppression has been found across species ^60–65^. Most notably, it has been studied in primates, where ascending pathways run from the midbrain ^6^ to the frontal eye field, a cortical structure involved in visual processing and oculomotor control ^66^. In rodents, the lateral posterior nucleus (LP), the homolog of the pulvinar nucleus, has also previously been implicated in correcting action-induced sensory signals, particularly saccades ^45,67^.

Beyond saccadic eye movements, reafference signals induced by self-motion will affect visual processing in a variety of ways. One approach would be for several different brain areas to act independently to generate CDs to compensate for different behaviors. However, it is challenging to maintain a reliable sensory representation through a distributed system. An alternative approach would be to pool all possible behavioral signals at a CD hub and modulate sensory signals according to the best overall estimate of the currently executed movement ensemble, ideally starting at the first possible step. Here, we provide evidence for the latter by showing that a strong inhibitory projection from the vLGN (Fig. 1) modulates visual signals in the superficial superior colliculus (Fig. 2). We found that this inhibitory projection mediates a combination of visual and behavioral parameters (Fig. 3) that are distinct from direct visual inputs from the retina (Fig. S3). In comparison, while retinal terminals show a variety of visual responses as previously shown ^40^, vLGN terminals are mainly driven by high-frequency luminance modulation and front-to-back visual motion (Fig. 3), stimuli that occur during forward locomotion in natural environments. Actions such as saccades, locomotion, or pupil dilation in visually homogeneous environments barely modulate retinal inputs, but strongly drive the vLGN projection. In addition, vLGN activity appears to relay direct copies of the motor action (Fig. 1 & 3), which would coincide with the arrival of visual inputs delayed by phototransduction. Accordingly, animals with a blocked vLGN showed saccade suppression deficits (Fig. 4), a computation that alleviates the visual blurring caused by different behaviors (Fig. 8A). Interestingly, such computations are achieved by largely preserving the temporal code of the first spike while reducing the total number of spikes (Figs. 2I-M and 4G-I), suggesting that the onset of visual signals remains informative and relevant irrespective if it is of reafferent or exafferent origin. While our data has been focused on the SC, anatomical evidence shows that other early sensory areas also receive direct inputs from the vLGN, e.g. the area LP and dLGN (Fig. 6), and thus are likely being similarly modulated. Consistently, vLGN blocked animals could not discriminate depth via motion parallax (Fig. 5). Given that the vLGN is an evolutionarily conserved structure ^20^, this CD motif is likely to be present across vertebrates. Taken together, our results reveal a powerful hub for sensory modulation that is a critical first step in attenuating the impact of sensory stimuli during action.

In the sensory periphery, the vLGN appears to regulate sensory signals, allowing the brain to compare predicted sensory inputs that originate through behavior with actual sensory signals. Such a feedback loop would enable constant adjustments of visuomotor transformations in real time ^1,6^, and thus improve perception. However, our functional and anatomical data suggest a more nuanced view beyond the classical view of the CD. The vLGN simultaneously coordinates and fine-tunes multiple sensorimotor processes. This is evident from optogenetic activation of the vLGN, which induces changes in locomotor speed, saccadic eye movements, and changes in pupil size (Fig. 6E-L). Accordingly, anatomical evidence shows that the vLGN projects and influences directly other subcortical target areas known to strongly influence some of these behaviors, such as the red nucleus and Edinger-Westphal nucleus via the olivary pretectal nucleus ^49^ (Fig. 6A-D, S6). Consistently, blocking vLGN output alters sensory evoked and internally controlled saccadic and pupil dilation dynamics (Fig. 7).

Action-induced sensory signals, such as motion blur, are generated by all kinds of actions, from locomotion to head and saccadic eye movements. Thus, a common correction in early sensory areas, i.e. SC, dLGN, LP (Fig. 1 & 6, Fig. S6), would be ideal. This is consistent with our results, which show that a mixture of behavioral features, in most cases in a bouton specific manner, are mapped to SC (Fig. 3I-M). However, behaviors can be highly variable in their kinetics and require specialized control systems, just think of a slow pupil dilation or a ballistic saccade. Thus, it is quite possible that the vLGN consists of a population of highly specialized feedback loops that operate in orchestration (Fig. 8B). This architecture facilitates both sensory adaptations that can be generalized across actions, such as motion blur compensation, and behavior-specific adaptations tailored to the dynamic properties of different actions. This notion is supported by single-cell sequencing data of the vLGN reporting a large neuronal diversity ^44^, previous findings that support divergent behaviors modulated by distinct vLGN projections ^15^, the diverse neuronal projections to areas related to specific behaviors (Fig. 6, Fig. S6), the diversity of behavioral patterns that emerge through optogenetic activation of the vLGN (Fig. 6E-L) and the behavioral deficits when the vLGN is blocked (Fig. 7). As there are some discrepancies in the literature regarding the exact projections of the vLGN ^16,29^, it is likely that these arise also from different neuronal populations with different projection patterns. Some will project mainly to the intermediate SC layers ^16^, others more to the superficial SC layers ^29^. In our study, we focused primarily on the superficial SC neurons, all of which receive strong inhibitory modulation from vLGN Gad2^+^ neurons (Fig. 1).

The vLGN is also known to relay visual information by receiving a variety of retinal inputs ^33^. However, all functional studies in the vLGN have lacked information on cell type specificity and thus, projection patterns. By functionally imaging the projections directly in the SC (Fig. 3), our results provide a valuable addition. Consistent with previous functional studies showing that the vLGN have large receptive fields that reflect changes in luminance ^29,41^. As previously shown, this information can refine and contextually modulate visuospatial processing, sharpening receptive fields and therefore supporting visually guided approach behavior ^29^ (Fig. 2). However, we also show that the strongest responses are due to high-frequency flicker and motion stimuli typically observed during forward translation (Fig. 3E-H). Such high-frequency and motion stimuli may occur when an animal runs under the canopy and may support other action cues conveyed by the vLGN. The exact computational role of such visuomotor interactions remains to be determined. Recently, the vLGN has also been implicated as an inhibitory switchboard for behavioral control ^68^ and, via its projections to the SC and nucleus reuniens, serves as a key regulator for adjusting defensive behaviors according to prior experience and the level of perceived visual threat ^15,16^. All of these findings suggest that CD signals are also modulated in a state-dependent manner, adding an additional layer of complexity to how animals process early sensory signals to adapt their perception to the demands of the moment.

In conclusion, the vLGN plays a fundamental role in the coordination of motor actions and visual processing, thereby maintaining perceptual stability. Our results underscore how closely vision and motion are linked to enable effective interaction with the sensory environment.

## LIMITATIONS OF THIS STUDY

In this study, we aimed to dissect the role of the vLGN in visuomotor functions by recording, stimulating and inactivating its neurons and tracing their input-output projections. The interpretation of these results is highly dependent on the specific targeting of vLGN neurons, for which we relied mainly on cell type-specific Cre recombinase expression in Gad2 neurons and targeted viral injections. Nevertheless, spillover into neighboring Gad2-expressing brain areas such as the IGL and the zona incerta (ZI) cannot be completely excluded and may contribute to the reported effects. We argue that the vLGN is the major contributor based on several lines of evidence: We verified qualitatively identical projection patterns between Gad2^+^ thalamic neurons (Fig. 1C-E) and V1-recipient thalamic neurons (Figs. 6A-C, S6, S7A-C) and showed that the majority of SC-targeting thalamic neurons are located in the vLGN (Fig. 1M,N), together ruling out a major contribution of the IGL. The ZI projects mostly to intermediate and deep layers of the SC ^69^, whereas our imaging of axonal terminals in the SC was limited to the superficial layers (Fig. 3). The position of the optogenetic stimulation fiber within the vLGN was verified histologically after the experiments (Fig. 2C), excluding stimulation in more distant areas, but likely including activation of IGL and minor lateral ZI neurons. Blocking vLGN transmission by TeLC expression could also affect surrounding areas (Fig. 4F). However, targeting TeLC expression medial to the vLGN (including ZI) did not affect visuomotor performance (Fig. S5G-J). Taken together, the vLGN appears to be the major contributor to the majority of the reported functions. Given the transcriptomic cell type diversity within the vLGN ^44^, it is likely that some of the wide range visuomotor functions reported here are distributed across several vLGN subcircuits. For example, we did not attempt to dissect whether visuomotor integration for depth perception relies on the same neurons and projections to the sSC that influence gaze shifting behavior. We hope that future studies will specify the diverse thalamic visuomotor functions to cell types and circuits within the vLGN.

## STAR METHODS

### RESOURCE AVAILABILITY

#### Lead contact

Further information and requests for resources and reagents should be directed to and will be fulfilled by the lead contact, Maximilian Jösch.

#### Materials availability

This study did not generate new unique reagents.

#### Data and code availability

Behavioral, *in vivo* electrophysiological, functional imaging data and *in vitro* electrophysiological patch clamp data will be deposited and made available as of the date of publication. DOIs will be listed in the key resources table. Microscopy data reported in this paper will be shared by the lead contact upon request.

All original code will be deposited and made publicly available as of the date of publication. DOIs will be listed in the key resources table.

Any additional information required to reanalyze the data reported in this paper is available from the lead contact upon request.

### EXPERIMENTAL MODEL AND SUBJECT DETAILS

#### Animals

Animal protocols were reviewed by the institutional preclinical core facility at IST Austria. All breeding and experimentation were performed under a license approved by the Austrian Federal Ministry of Science and Research in accordance with the Austrian and EU animal laws (BMF-66.018/0017-WF/V/3b/2017). During the experimental phase, mice were housed individually in standard macrolon cages with red plastic houses, running wheels and enrichment consisting of wood chips and nesting material at an inverted 12-hour light cycle. Experiments were done during the dark phase of the light cycle.

For *in vivo* tracing (n = 10, 6 males (m), 4 females (f)), *ex vivo* patch-clamp (n = 8, 5 m, 3 f), *in vivo* opto/electrophysiology (n = 6, 4 m, 2 f), *in vivo* vLGN terminal imaging (n = 6, 4 m, 2 f), and *in vivo* TeLC experiments (n = 12, 6 m, 6 f), Gad2-IRES-Cre (JAX, #010802), aged 8 weeks (5–12 weeks for vLGN bouton imaging) at viral injection were used. For *in vivo* anterograde transsynaptic experiments, Ai75D (JAX, # 025106, n = 6, 4 m, 2 f), aged 8 weeks at viral injection were used. For *in vivo* retrograde transsynaptic experiments, NTSR1-GN209-Cre (MMRRC, #030780, n = 4, 2 m, 2 f), GRP-KH288-cre (MMRRC, #031183, n = 4, 2 m, 2 f), and Rorb-Cre (JAX, # 023526, n = 4, 2 m, 2 f), aged 8 weeks at viral injection were used. The mice for retinal terminal imaging experiments were C57BL/6J (JAX, #000664; n = 5, 3 m, 2 f), aged 6–11 weeks at eye injection. Of those, 3 mice have been used to record previously published separate datasets ^42^.

### METHOD DETAILS

#### Statistics and Reproducibility

No statistical method was used to predetermine sample size. Extracellularly recorded units and imaged bouton ROIs were selected or excluded based on quality and response criteria as described below.

#### Viral vectors

Anterograde transsynaptic expression was done with AAV1-cre (AAV1.CamKII0.4.Cre.SV40, 7 × 10¹² genome copies per ml, Addgene). Retrograde transsynaptic expression was performed with starter vector (AAV-DIO-Ef1a-TVA-FLAG-2A-N2C_G) ^30^ and pseudotyped rabies vector (N2C(Enva)-EGFP, ∼2–5 x10^8^ genome copies per ml) ^30^. Calcium indicator expression in vLGN/IGL neurons was achieved with AAV-hSynapsin1-FLEx-axon-GCaMP6s (1 × 10^12^ genome copies per ml, Addgene, 112010-AAV5) for retinal expression AAV2.7M8-syn-GCaMP8m viral vectors (1 × 10^13^ genome copies per ml), generated at ISTA viral facility. Tetanus Neurotoxin Light Chain (TeLC) viruses were generated using AAV5-hSyn-FLEX-TeLC-P2A-dTomato (Addgene, 159102, 1 × 10^13^ genome copies per ml) at ISTA viral facility. ChR2 (AAV5-EF1a-doubleFloxed-hChR2(H134R)-EYFP-WPRE-HGHpa (#20298-AAV5), 1×10¹³ genome copies per ml), mCherry control (AAV5-hSyn-DIO-mCherry (#50459-AAV5), 7 × 10¹² genome copies per ml) and eNpHR3.0 (AAV5-hSyn-eNpHR3.0-EYFP (#26972-AAV5), 10^¹3^ genome copies per ml) viruses were purchased from Addgene.

#### Stereotaxic viral injections

Anesthesia was induced with 3 % isoflurane and intraperitoneal (i.p.) ketamine and xylazine (100 mg kg^−1^, 10 mg kg^−1^). As an analgesic, meloxicam (20 mg kg^−1^) was subcutaneously (s.c.) injected. Mice were placed in a stereotaxic apparatus (Kopf) and body temperature was controlled with a heating pad at 37 °C throughout the whole procedure. Stereotaxic target coordinates relative to bregma were −2.3 mm (AP), 2.5 mm (ML), 3.4 mm (DV) for vLGN, - 2.3 mm (AP), 2.3 mm (ML), 3.4 (DV) for medial thalamus, −3.8 mm (AP), 0.8 mm (ML), 1.3 mm (DV), for SC and 4 mm (AP), 2.6 mm (ML), 0.5 mm (DV) for visual cortex injection. Glass electrodes were pulled with a one-stage puller (DMZ-Zeitz-Puller) to produce a tip opening ∼30 μm. The pipette was filled with mineral oil, then attached to a Nanoliter 2010 (World Precision Instruments) and loaded with the respective vector. Pipettes were slowly lowered to the target region (vLGN 150nL / 300nL, and SC 200 / 300 nl, V1 60 nL) and the solution was injected at a rate of 45 nl min^−1^. Once the volume was delivered, pipettes remained in place for 15 min before being carefully withdrawn and the incision closed with VetBond (3M). For retrograde transsynaptic tracing, the pseudotyped rabies vector was injected 7 days after the first starter vector injection. Otherwise, animals were recovering and awaiting viral expression for at least 3 weeks, before further experiments were conducted. For vLGN terminal imaging experiments, in 4 out of 6 mice, viral infection was immediately followed by cranial window implantation in the same surgery. The remaining 2 mice were implanted 4 weeks after the injection surgery.

#### *In vitro* electrophysiology

Mice were deeply anesthetized via intraperitoneal (i.p.) injection of ketamine (95 mg kg^−1^) and xylazine (4.5 mg kg^−1^), followed by transcardial perfusion with ice-cold, oxygenated (95 % O_2_, 5 % CO_2_) artificial cerebrospinal fluid (ACSF) containing (in mM): 118 NaCl, 2.5 KCl, 1.25 NaH_2_PO_4_, 1.5 MgSO_4_, 1 CaCl_2_, 10 Glucose, 3 Myo-inositol, 30 Sucrose, 30 NaHCO_3_; pH = 7.4. The brain was rapidly excised and coronal sections of 300 µm thickness containing the SC were cut using a Linear-Pro7 vibratome (Dosaka, Japan). Slices were left to recover for 20 min at 35 °C, followed by a slow cool down to room temperature (RT) over 40 – 60 min. After recovery, one slice was transferred to the recording chamber (RC-26GLP, Warner Instruments, Holliston, MA, USA) and superfused with ACSF containing 2 mM CaCl_2_ at a rate of 3 – 4 ml/min at RT (21.0 - 23.0 °C). Glass pipettes (B150-86-10, Sutter Instrument, Novato, CA, USA) with resistances of 3 – 4 MΩ were crafted using a P1000 horizontal pipette puller (Sutter Instrument) and filled with internal solution containing (in mM): 140 K-Gluconate, 2 MgCl_2_, 2 MgATP, 0.2 NaGTP, 0.5 EGTA, 10 HEPES; pH 7.4 adjusted with KOH. Biocytin (0.2 – 0.3 %) was added to the internal solution for post hoc morphological reconstruction. Electrical signals were acquired at 20 – 50 kHz and filtered at 4 kHz using a Multiclamp 700B amplifier (Molecular Devices, San Jose, CA, USA) connected to a Digidata 1440A digitizer (Molecular Devices) with pClamp10 software (Molecular Devices). For optogenetically evoked inhibitory postsynaptic currents (IPSCs), neurons were held at −60 mV and blue light (λ = 465 nm, 10 – 20 mW/cm^2^ intensity, 5 ms pulse duration, 0.1 - 0.2 Hz stimulation frequency) was emitted through a mono fiber-optic cannula (5 mm length, fiber diameter 200 μm, total diameter 230 μm, Doric lenses, Quebec, Canada) connected to a PlexBright LED 644 (Plexon, Dallas, TX, USA) with an optical patch cable (fiber diameter 200 μm, total diameter 230 μm, 0.48 NA). To block GABA_A_ receptors, ACSF containing 20 µM bicuculline was bath-applied for 20-30 s followed by immediate washout. Access resistance was constantly monitored between protocols and recordings with access resistances exceeding 20 MΩ or with changes in access resistance or holding current by more than 20 % were discarded. After recordings, the pipette was carefully withdrawn and the slice was transferred to 4 % paraformaldehyde (PFA) in PBS solution.

#### Viral eye injections

For expression of calcium indicators in retinal neurons, C57BL/6J mice were anesthetized with ketamine/xylazine (100 mg kg^−1^, 10 mg kg^−1^) by i.p. injection. A small hole in the temporal eye, below the cornea, was cut with a 1/2-inch, 30-gauge needle. Subsequently, 1 μl of vitreous fluid was withdrawn and 1 μl of AAV2.7M8-syn-GCaMP8m viral vector solution was injected into the subretinal space with a Hamilton syringe and a 33-gauge blunt-ended needle. Mice were left to recover and viral expression was to commence for 2-4 weeks before implantation of the cranial window.

#### Cranial window implantation surgery

Cranial window implantation was performed as described previously ^42^. In brief, mice were injected with meloxicam (20 mg per kg body weight, s.c., 3.125 mg ml^−1^ solution) and dexamethasone (0.2 mg per kg body weight i.p., 0.02 mg ml^−1^ solution). Anesthesia was induced by 2.5 % isoflurane in oxygen in an anesthesia chamber and maintained at 0.7 % to 1.2 % in a stereotaxic device (Kopf), while body temperature was controlled by a heating pad to 37.5 °C. After exposing and cleaning the cranium, a 4 mm circular craniotomy was drilled above the left SC, the dura mater was removed, the left transverse sinus was sutured twice with 9-0 monofil surgical suture material (B. Braun) and cut between the sutures. Cortical areas covering the left SC were aspirated with a cell culture vacuum pump (Accuris), and a 3 mm circular coverslip, glued (Norland optical adhesives 61) to a stainless-steel conical ring, was inserted with the glass flush on the surface of the SC. After filling the surrounding cavity with Dura-Gel (Cambridge Neurotech) the insert was fixed in place with VetBond (3M). Finally, a custom-designed TiAl_6_V_4_ head-plate was affixed to the cranium by sequential application and curing of (1) All-in-One Optibond (Kerr), (2) Charisma Flow (Kulzer), and (3) Paladur (Kulzer). Mice were given 300 µl of saline and 20 mg per kg body weight meloxicam (s.c.) before removing them from the stereotaxic frame and letting them wake up while keeping them warm on a heating pad. Another dose of 20 mg per kg body weight meloxicam s.c. and 0.2 mg per kg body weight i.p. dexamethasone was further injected 24 h after the conclusion of the surgery. After the implantation surgery, mice were allowed to recover for at least one week.

#### Setups for head-fixed *in vivo* recordings

For awake behaving experiments, two similar setups were used, with the difference that one was coupled to a custom-built multiphoton setup, and the other allowed for silicon probe/neuropixels recordings. In short, mice were head-fixed while awake using a custom-manufactured clamp (for imaging: connected to a three-axis motorized stage (8MT167-25LS, Standa)) and could run freely on a custom-designed spherical treadmill (20-cm diameter). Running behavior was recorded by a pair of ADNS-3080 (iHaospace, Amazon) optical flow sensor modules, focused with 25 mm lenses (AC127-025-AB-ML, Thorlabs) on a small patch at orthogonal locations of the Styrofoam ball and illuminated by an 850 nm LED. The alternating sensor readout was controlled at 50 frames/s by an Arduino Uno running custom scripts. The 4 signal channels from the sensor were linearly mapped to movement speed in the forward, sideways and rotational axes based on regular calibration with synchronous measurement of image translations and rotation at the ball’s apex. Eye and body movements were recorded at 50 fps with infrared illumination (850 nm) with a Camera (acA1920-150um, Basler) and an 18-108 mm macro zoom objective (MVL7000, Thorlabs) for multiphoton imaging or a fixed focal length objective for electrophysiology (Edmund Optics, f = 50 mm #59-873), pointed at the right side of the mouse via an infrared mirror. Eye position and saccades were determined post-hoc as previously reported ^42^, by first labeling eight points around the pupil with DeepLabCut ^70^, which were fitted to an ellipse, and the center position was transformed to rotational coordinates. Fast eye position changes of more than 45 °s^−1^ and at least 3 ° amplitude on a 0.7 s median filtered trace were defined as saccades. The ellipse area in mm^2^ was determined as pupil size.

Visual stimuli were projected by a modified LightCrafter (Texas Instruments) at 60 Hz (Multiphoton setup: DLP Lightcrafter evaluation module; e-phys setup: DLP LightCrafter 4500, Texas Instruments), reflected by a quarter-sphere mirror (Modulor) below the mouse and presented on a custom-made spherical dome (80 cm in diameter) with the mouse’s head at its center. For imaging experiments, a double bandpass filter (387/480 HD Dualband Filter, Semrock) was positioned in front of the projector to minimize light contamination during imaging. In both setups, the blue LED in the projector was replaced by UV (LZ1-00UB00-01U6, Osram) and in addition, in the multi-photon setup, the green LED was replaced by a cyan LED (LZ1-00DB00-0100, Osram) not to interfere with the Calcium imaging wavelengths. The reflected red channel of the projector was used for synchronization and captured by a trans-impedance photo-amplifier (PDA36A2, Thorlabs) and digitized. Stimuli were designed and presented with Psychtoolbox-3 ^71^, running on MATLAB (MathWorks) on Microsoft Windows 10 systems. Stimulus frames were morphed on the GPU using a customized projection map and an OpenGL shader to counteract the distortions resulting from the spherical mirror and dome. In both setups, the dome allows the presentation of mesopic stimuli from circa 100 ° on the left to ca. 135 ° on the right in azimuth and from circa 50 ° below to circa 50 ° above the equator in elevation. In between dynamic stimuli presented in randomized order, the screen was set to a homogeneous gray (green & UV light) at scotopic level for at least 30 s. To determine behavioral coupling, these stimuli were interspersed with 5 min gray screens, i.e., at visual baseline.

#### *In vivo* electrophysiology and optogenetics

Gad2-Cre mice, previously injected with AAV5-EF1a-doubleFloxed-hChR2(H134R)-EYFP were anesthetized with isoflurane (1 % - 1.5 % in oxygen 0.8 l min^−1^) and injected with meloxicam (20 mg kg^−1^, s.c.) and placed in the stereotaxic apparatus. The skull was exposed and the periosteum and connective tissue removed. Thin crossed grooves over the bone were cut to increase the contact surface using a scalpel. The skull was first covered with a thin layer of cyanoacrylate (VetBond, 3M), then Charisma Flow (Kulzer) that was blue-light-cured for 45 s, before securing a head plate with Super-Bond dental adhesive resin cement (Sun medical). A tapered optic **λ**-fiber with an active zone of 0.5 mm (NA 0.39, Optogenix) was implanted using the same vLGN coordinates and craniotomy as the injection. The tip of the fiber was slowly lowered to a depth of 3.4 mm from the dorsal surface and cemented to the skull.

One day prior to the recording session, mice were anesthetized with isoflurane (1 % - 1.5 % in oxygen 0.8 l min^−1^) and injected with meloxicam (20 mg kg^−1^, s.c.). A small craniotomy was made in the rostral skull (Bregma: 0.5 mm AP, 2 mm ML) for implanting an inverted gold pin as a reference electrode. A second rectangular craniotomy was made over the Cortex/SC region (Bregma: −3.5-3.8 mm AP, 0.5-1 mm ML), leaving the dura mater intact. The window was covered with silicone elastomer (Kwik-Cast, World Precision Instruments). The next day, Kwik-Cast was removed and the well around the craniotomy was constantly filled with ACSF throughout the whole recording session. Extracellular recordings were obtained using a single shank acute linear 32-channel silicon probe (ASSY-37 H4 with probe tip sharpening, Cambridge Neurotech) connected to an RHD 32-channel amplifier board and RHD2000 USB Interface Board (Intan Technologies) and Neuropixels 2.0 multishank probes (IMEC), usign a Neuropixels data-aquisition system (see Neuropixels.org for more detail). Before recording, the tip of the electrode was coated with DiI (Invitrogen) to allow post hoc recording site location. To access the superficial SC, the probe was slowly inserted through the cortex at a speed of 1 μm sec^−1^ to a depth of ∼ 1.7 mm using a stable micromanipulator (Luigs & Neumann Motorized, Germany). The electrode was left in place for 30 min before starting to record. Data were sampled at 20 kHz using Labview 2017 (National Instruments). Spike-sorting was performed with Kilosort 2 (https://github.com/cortex-lab/Kilosort) ^72^. The automatic template of Kilosort 2 was manually curated on Phy2. The 473 nm laser (SDL-473-XXXSFL-RA, Shanghai Dream Laser Technology Co. Ltd) bursts for optogenetics were generated in Arduino Due (www.arduino.cc) in pulses of 40 Hz with an approximate power at the fiber tip of 2.5 mW/mm^2^.

#### Visual local flash and optogenetics

Prior to starting the experiment, the visual field was scanned with a dark disk with 10° radius to determine the approximate location of receptive fields. We used this location to present a white or dark disk of the same radius. The visual local flash was interleaved in time with the laser burst of the same length, stimulating optogenetically vLGN. The duration of the laser and visual stimulation was 200 ms (Fig. 2E-G, I) or 1 sec (Fig. 2H, Fig. S8A-F). Laser burst offsets relative to the start of the visual flash were varied in 13 randomized order increments with the start of the burst *i* set to −1.5*t_flash_ + *i*\* t_flash_/4 for *i* in [0,12], where t_flash_ was the duration of the flash.

### QUANTIFICATION AND STATISTICAL ANALYSIS

#### Current Source Density Analysis

To confirm the location of the silicone probe during the *in vivo* recordings, current source density analysis ^73,74^ was applied. For this analysis, local flash stimuli that were at least 0.5 s after the laser burst were used. For each such repetition of the flash, CSD profile ^74^ was computed on the raw voltage recorded values in the interval [−0.1, 0.2] s around the flash onset and averaged over multiple repetitions. The channel of the silicone probe corresponding to the current sink is defined as the channel where the current flow is the smallest. The closest channel above with positive current flow is the source. The depth of the source channel was set to 300 μm; the depth of the remaining channels was derived relative to the source using the 25 μm spacing between the channels. The response magnitude (Fig. 2F) was computed as the variance of the CSD profile of each channel across time. The normalized response (Fig. 2G) is the variance across all channels after the variance before the onset of the flash was subtracted and normalized to the maximum. The same procedure was applied to compute the CSD analyses around laser bursts but selecting laser burst onsets that were at least 0.5 s after a visual flash.

#### Neuronal responses

Both the zeta-test ^75^ and a permutation test ^76^ with subsampling were used to identify units that were responsive (p < 0.01) to the visual or optogenetic stimulation. The two tests detect complimentary response types: the zeta-test identifies event-locked responses, whereas the permutation test captures changes in the mean firing rate, including tonic changes of the firing rates, such as in the case of optogenetic stimulation of vLGN/IGL complex. For the permutation tests, the firing rate during the stimulus was compared to the baseline firing rate, estimated from random samples of 0.2 s intervals before the stimulus. Units were defined as *visually responsive* if they had either ON or OFF responses to the flash within 0.2 s after the start/end of the flash. A unit was considered *optogenetically responsive* if it exhibits a change of the spontaneous firing rate during 0.2 s after the start or end of the laser burst or if its visual ON/OFF responses were altered in the presence of optogenetic stimulation (permutation test). For analysis in Fig. 2H-L only units responsive to both the visual flash and optogenetic stimulation were selected. To compute responses to visual flashes, optogenetic stimulation, and both of the above (Fig. 2H-J and S2B-D,F) we used the trials where the visual flash preceded optogenetic stimulation (visual responses, Fig. 2H, S2B left), or vice versa (optogenetic responses, Fig. 2H, S2B middle), or where visual and optogenetic stimulation overlapped (visual & optogenetic responses, Fig. 2H, S2B right). In Fig. 2L, S2E, the mean response of all units per laser offset was computed after normalizing the responses of each unit to their maximum across all laser offsets. To analyze receptive fields (RF, Fig. 2M,N) a vertical bar (size 2° and 8°) was presented at a random horizontal position on the screen, the location of the bar was updated with the frequency 15Hz. At the same time the pulses of optogenetic stimulation of LGN/IGL complex happened with the frequency of 1.4Hz and pulse duration of 0.1 s, this short duration of optogenetic pulses did not cause pupil dilation. To avoid the effect of rebound spiking, the spikes within 0.1 s after optogenetic pulses were removed from the analysis. We subsampled the spikes so that the number of spikes in conditions with and without optogenetic stimulation match for each unit. To reconstruct RFs, we averaged the frames presented during [−0.5,0.08] s around each spike, separating conditions into groups with and without optogenetic stimulation at the time of the spike. For further analysis, we excluded the units with noisy RF in either of the two conditions. For this, signal to noise ratio (SNR) was computed as the ratio of the variance of the RF in the time interval [T-1, T+1], to the variance outside of this interval, where T is the time of the maximal RF variance. The threshold for SNR was set to 80th percentile of SNR of all units, at the value 3.44. The horizontal profile of a RF (Fig. 2M bottom, black) was computed as the mean of three frames around T. We fit the 1D Gaussian function (Fig. 2M bottom, blue) g(x)=A*exp((x-m)^2^/(2w^2^))+b, where the parameters A, m, w, and b are the amplitude, mean, width, and the baseline. The fitting was done using lsqcurvefit Matlab function. The width of the RF (Fig. 5N) was estimated using the fitted parameter w.

#### Optogenetically triggered behavior analysis

For vLGN, tapered optic **λ**-fibers with an active zone of 0.5 mm (NA 0.39, Optogenix) were implanted using the same vLGN coordinates as vector injections. For SC, optic fibers (400 μm diameter, NA 0.39, ThorLabs) were implanted at 1000 μm from the pial surface using the same AP and ML coordinates as vector injections. Both types of fiber were fixed using light-curing glue (Optibond Universal, Kerr Dental) and dental cement (SuperBond C&B Kit, Hentschel-Dental).

To analyze optogenetically triggered behaviors, only 1 s optogenetic stimulation pulses, where the offset preceded the visual flash, were included. To determine turning speed (Fig. 6K), mean speed in a window of 0.25 s before stimulation onset was subtracted and trials sorted by mean speed in a 0.5 s window after stimulation onset, for visualization. Maximum turning speed (Fig. 6L) is the maximum within 0.25 s after the laser onset. Change of azimuthal pupil position (Δ az, was normalized and sorted using the same windows, defining starting position as the mean pupil azimuth between 0.25 s before and laser onset (Fig. S8B,D,F left). Pupil velocity (Fig. S8A,C,E) was computed as a central difference of sequential pupil azimuth values *v_t_ = (az_t+1_ - az_t-1_)/2.* Maximum pupil velocity (Fig. 6H) is the maximum of the velocity profile of each trial in the 0.25 s after the laser onset, the mean of each group was computed from the trials in Fig. S8A,C,E). Change in pupil azimuth (Fig. 6G) was defined as Δ*az = az_T-w/2_ - az_T+w/2_*, with the location *T* and width *w* of the maximum peak in the pupil velocity profile computed using the Matlab function findpeaks. To determine changes of pupil size (Fig. 6I-J, Fig. S8H,J), mean pupil diameter was computed during 0.25 s before and after laser offset; the difference between the two conditions was estimated using the Wilcoxon signed rank test (p = 10^−74^). Fig. 6J shows the means for the pupil changes for the three conditions Fig. 6I, Fig. S8H, and Fig. S8J.

#### *In vivo* vLGN terminals imaging

Two-photon terminal imaging in superior colliculus was performed as described previously ^42^. In brief, ScanImage (Vidrio Technologies) on MATLAB 2020b (MathWorks) controlled a custom-built microscope using a pulsed Ti:Sapphire laser (Mai-Tai DeepSee, Spectra-Physics) set at wavelengths between 920 and 950 nm. The beam was expanded to underfill the back-aperture of the objective (×16 0.8-NA water-immersion, Nikon) and scanned through the tissue by a galvanometric-resonant (8 kHz) mirror combination (Cambridge Scientific) and a piezo actuator (P-725.4CA, Physik Instrumente) controlling the objective. Emission light was measured with GaAsP photomultiplier tubes (H10770B-40, Hamamatsu) following collection by a dichroic mirror (FF775-Di01, Semrock) and channel splitting (580 nm long-pass, FF580-FDi01, Semrock) as well as filtering (green: FF03-525/50; red: FF01-641/75; Semrock). The signals were then amplified by a TIA60 amplifier (Thorlabs) and digitized with a PXI system (PXIe-7961R NI FlexRIO FPGA, NI 5734 16-bit, National Instruments). Average laser output power at the objective ranged from 38 to 125 mW (median of 75 mW). A FOV of 0.13– 1.85 mm^2^ (median of 0.77 mm^2^) was imaged over 3–12 planes (median of 6 planes) with a plane distance of 10–45 µm (median of 28 µm) at a pixel size of 0.6–1.9 µm (median of 1.3 µm) and a volume rate of 4.2–9.5 Hz (median of 4.8 Hz). The field of view varied between recordings, ranging from 0.2 to 1.8 mm^2^ (median = 0.7 mm^2^) of the SC surface for vLGN/IGL terminal imaging and 0.1 to 1.6 mm^2^ (median = 0.7 mm^2^) for retinal bouton imaging. Each mouse was recorded in 1-9 (median of 7) imaging sessions on different days. In a subset of recordings (n = 15) in separate imaging sessions, the absence of substantial z-motion was verified by injecting 40 µl of Texas Red dextran (3000 MW, 14.3 mg ml^−1^, diluted in saline, Thermo Fisher Scientific) s.c. and imaging brightly red labeled blood vessels at 980 nm ^77^.

#### Visual stimuli for *in vivo* terminals response mapping

To measure sensitivity to luminance dynamics, repeated sequences of luminance chirps, as reported previously^42^, were used. The stimulus started at gray level, followed by a 1 s bright step and sinusoidal luminance changes over 8 seconds each, first with increasing amplitude at 2 Hz and then fixed full amplitude but frequency modulated (0 to 8 Hz). For determining direction selectivity, sinusoidal gratings of 0.1 cycles per degree spatial frequency, and 2 cycles per second temporal frequency were presented at full contrast moving in 8 or 16 directions in randomized order. Gratings were presented stationarily for 3 s and then moved for 7 s in the current direction. To test for retinotopy, a dark bar with length spanning the screen and width of 25 ° was moved over gray background at 22.5 °/s in 8 directions perpendicular to the bar orientation for 7 s with 3 s interval between presentations. Full field flash responses were determined by presenting either dark or white 1 s full field flashes from gray baseline at a pseudorandom interval of 5 to 10 s. Pseudosaccade stimuli consisted of vertical gratings with 0.08 to 0.25 cpd spatial frequency, or random checkerboard patterns with 4 to 12 ° visual angle checker size, which were presented on the screen. At pseudorandom intervals between 3 to 6 s the full screen texture moved in a random horizontal direction over 0.08 s by a median of 5 ° (3 to 30 °) visual angle. The distribution of such pseudosaccadic image shifts approximately matched those from actual saccades, as determined from head-fixed population spontaneous saccade statistics.

#### *In vivo* axonal terminal imaging analysis

Imaging data was motion corrected and ROI segmented with suite2p (v0.10.0)^78^ followed by a manual curation step based on morphological and activity shape. Note that multiple axonal ROIs can originate from the same neuron. Further analysis was performed in MATLAB (MathWorks). dF/F_0_ was estimated as previously ^42,79^, by subtracting neuropil contamination with a factor of 0.5, defining F_0_ baseline as the 8th percentile of a moving window of 15 s ^80^ and finally subtracting and then dividing the fluorescence trace by the median of the same 15 s window. The fluorescence signal-to-noise ratio was defined for each ROI by dividing the 99^th^ percentile of the dF/F_0_ trace (‘signal’) by the standard deviation of its negative values after baseline correction (‘noise’). Only axonal segments with a fluorescence SNR ≥ 5 were included in further analysis.

To estimate modulation by visual and behavioral stimuli, a suite of stimuli (full field flashes, frequency modulation, moving bar, moving full field grating) was presented and a range of behaviors (locomotion, pupil size, saccades) were sampled. Modulation indices were computed by (F_response_ - F_baseline_) / (F_response_ + F_baseline_), F being the average dF/F_0_ value in a baseline or response window, respectively. For moving bar and grating stimuli, baseline window was defined as [1.5, 0.1] s before movement start, response window from 0.25 after stimulus start until the end of the stimulus. For full field luminance modulation baseline was between 1.3 s before and until frequency modulation start, response as the time of frequency modulation presentation. For 0.5 s black or white full field flashes, baseline window was 1 s before flash onset, response window was 1 s following flash onset, including OFF responses. For saccades and locomotion onset [1.5, 0.5] s before were used as baseline and [0, 1] s following onset was used as response window. Significance of modulation was determined by two-sided Wilcoxon signed-rank tests. Significance of correlations was determined by 5000 repeats of randomly shifting the behavioral trace between 120 s and 1200 s and computing correlations for shuffled datasets. Significance was then determined by calculating the proportion of shuffles with more extreme correlation values than the actual data. Boutons from either vLGN or retina were included in further analysis if they showed Bonferroni-corrected significant (p < 0.01) modulation or correlation to at least one out of 8 tested conditions. Note that not all recordings included all stimulus sections. These inclusion criteria removed 102833/322854 vLGN and 22646/101376 RGC over all recorded boutons from further analysis (Figs. 3, 5E-H, S3, S4).

For pseudosaccade analyses (Fig. 5E-H), baseline window was defined as [0.5, 0.1] s before and response window as [0, 0.5] s following the pseudosaccade or saccade. Only saccadic events separated by at least 0.75 s from other saccadic events were included to avoid cross-contamination. Population synchrony (Fig. S4A) was determined as variance of the population mean divided by the population mean of individual bouton variance. Similarly, individual bouton variance explained (Fig. S4B) was determined by z-scoring bouton activity, and computing remaining variance after subtracting population mean (var_exp_ = 1 - var[act - <act>_population_]_t_, with act being z-scored bouton activity). To determine direction selectivity and preferred directions, 1 - circular variance and vector sums were used^81^. To demonstrate preferred direction distribution (Figs. 3F and S3F), only boutons with significant direction tuning p < 0.01 (10000-fold shuffled direction label test) were plotted. To plot retinotopic alignment of bouton responses (Figs. 3G and S3G), mean bouton responses to dark bars moving in nasal-temporal direction were determined. The centroids were projected into a set of 1-D axes, rotated at angles from 0° to 180° with the increment of 5°, and binned at 20 µm. The responses of boutons within each bin were averaged. The axis that yielded the maximal correlation of the binned response peak latency with the horizontal position of the bar was used for the alignment and the corresponding binned responses are shown in Figs. 3G and S3G. To determine cross-correlation timing (Fig. S4D,E), lag time of cross-correlation maximum was determined and bouton included if peak lag was within [−3, 3] s and p < 0.01 (random shift test, see above). To disentangle independent locomotion and pupil size contributions to bouton correlations (Fig. S4F-I), correlations to pupil size were separately computed for stationary periods (0.25 s window median filtered locomotion speed < 1 cm/s). Due to large sample sizes, comparisons between bouton populations yield arbitrarily low p-values. In these cases, mean and standard deviation of the difference are reported alongside.

#### Visual cliff setup

The visual cliff paradigm was performed in a black walled 50 x 50 cm acrylic box with 80 cm height, covered with transparent (5 mm thickness) and surrounded by black acrylic walls (∼25 cm height). The illusory platform (25 x 25 cm) was created by gluing a paper-printed checkerboard pattern to the bottom and adding a matching black acrylic border (W: 1cm x H: 0,5 cm) to the surface of the transparent acrylic surface. Interior walls beneath the transparent surface as well as the box floor were covered with a high-contrast checkerboard pattern (2.5 x 2.5 cm sized black and white squares) so that all edges were aligned. One day before experiments, mice were shortly anesthetized with isoflurane (5 %) and vibrissae clipped next to the mystacial pad with surgical scissors. Once recovered, mice were returned to the maintenance cage. A camera (Basler acA1920-150um) with a fixed focal length objective (Edmund Optics, f = 50 mm #59-873) was located above the middle point of the arena to cover all movements. Camera control and recordings were obtained using a custom-made script in Python. Each mouse was recorded for 30 min while freely roaming in the arena, only the first 10 minutes of which are included in further analysis.

#### Visual cliff analysis

The head and body were tracked using a custom-trained network in DeepLabCut ^70^. Ear tag labels were used for trajectory analysis as they were the most reliable. Video frames were cropped to the arena size and scaled to 1000 x 1000 pixels for consistency. *Platform area* was defined as the bottom left 25 cm quadrant of the arena, the remainder as *cliff area*. The cliff avoidance index AI (Fig. 5D,E and Fig. S5H,I) was computed as *AI = (t_platform_ - t_cliff_) / (t_platform_ + t_cliff_)*, where t is the time spent in the platform or cliff normalized by the area of these regions. To compute the avoidance index, a 10 cm strip along the walls was excluded (Fig. 5B). Aborted exits (Fig. 5F,G and Fig. S5J) were counted when a mouse crossed the platform boundary from inside the platform but then reversed the direction of movement normal to the platform boundary. Aborted exit trajectories were extracted for 4 s around these timepoints and multiple aborted exits removed.

#### vLGN inactivation physiology

To quantify responses to saccades (Fig. 4G-I), the recordings during oscillating random checker stimulus were used. A random checker pattern (8° per checker) was oscillating sinusoidally by 17 ° in the horizontal direction for 450 cycles per recording, analogous to previous reports ^53^. Up to two sessions were recorded per mouse, however, due to synchronization problems, some recordings had to be discarded. Only the first available recording per animal was used for the analysis. Saccades were identified as described above. Zeta-test ^75^ was used to idendify saccade-responsive neurons. To test optokinetic reflexes (Fig. 7A-E), the same stimulus was used. The starting pupil position per cycle was subtracted from the pupil position traces (Fig. 7C). Saccadic events (Fig. 7D-E) were identified as pupil displacements above 2.5 °/frame. To determine luminance change responses, 1 s full-field flashes of three different intensities (low, medium, and high) were presented in a random order. Pupil size response speed (Fig. 7G-H) was determined by estimating the slope of a linear fit to the relative pupil area change (baseline subtracted, fitted to the pupil values 0.2 s after flash onset). Quiescent and locomoting states (Fig. 7I-K) were identified as forward locomoting speed below and above 5 cm/s. Zeta-test ^75^ was done to detect visually responsive neurons, and PSTH for white and black flashes was computed (Fig. S5D-I).

#### Euthanasia and histology

Mice were dosed with 750-1000 mg kg^−1^ mixture of ketamine/xylazine and transcardially perfused with PBS, followed by ice-cold 4 % paraformaldehyde (PFA) in PBS. Brains were carefully extracted and post-fixed overnight, cryoprotected with sucrose 30 %, and sectioned at 60 μm using a sliding microtome (Leica SM2010). Sections were collected in three series. The first series was used for signal amplification of the respective vector. Briefly, sections were incubated with PBST (Triton 0.3 %) solution containing 5 % donkey normal serum and one or more antibodies (goat Anti GFP, ab6673, Abcam, diluted 1:2000; rabbit anti-RFP, 600-401-379, Rockland, diluted 1:1000) overnight at 4 °C, followed by secondary fluorescent antibodies (Donkey anti-goat-488, ab150129, Abcam, diluted 1:1000; Donkey anti-rabbit-594, R37119, ThermoFischer, diluted 1:1000) at room temperature for 1 h. Sections were mounted on slides and coverslips with custom-made mowiol.

#### Confocal microscopy

Brain sections were imaged with a Nikon CSU-W1 spinning disk confocal microscope. All images were processed with FIJI (ImageJ).

#### Statistics

Analyses were performed in custom-written MATLAB (MathWorks, Inc.) and Python scripts. Non-parametric tests used and defined in the figure legends. All statistical tests are reported in the text and appropriate figure legends (\**p* < 0.05, \*\**p* < 0.01, \*\*\**p* < 0.001). In bar plots the mean ± s.e.m. are shown, unless otherwise stated.

## ACKNOWLEDGEMENTS

We thank Yoav Ben Simon for generously making viral vectors for retrograde tracing available, as well as Jake Watson and Florian Marr for reagents. We also thank Ryuichi Shigemoto, Wiktor Młynarski and members of the Neuroethology group for their comments on the manuscript. This research was supported by the Scientific Service Units of IST Austria through resources provided by Scientific Computing, the Preclinical Facility, the Lab Support Facility, and the Imaging and Optics Facility, in particular Freyja Lange, Michael Schunn and Todor Asenov. This work was supported by European Research Council Starting Grant 756502 & European Research Council Consolidator Grant 101086580 (MJ) and EMBO ALTF 1098-2017 (AS) Human Frontiers Science Program LT000256/2018-L (AS).

## AUTHOR CONTRIBUTION

TVZ, AS and MJ designed the study. TVZ performed the *in vivo* electrophysiology, anatomical characterization and behavioral experiments. PK performed the slice physiology. AS performed the *in vivo* imaging experiments with help from FS. TVZ, AS, MJ and OS analyzed the experimental data. Software engineering for hardware control was performed by OS and AS. TVZ, AS and MJ wrote the manuscript with help of the other authors.

