## Supplementary Figures for "A thalamic hub-and-spoke circuit enables visual perception during action by coordinating visuomotor dynamics"

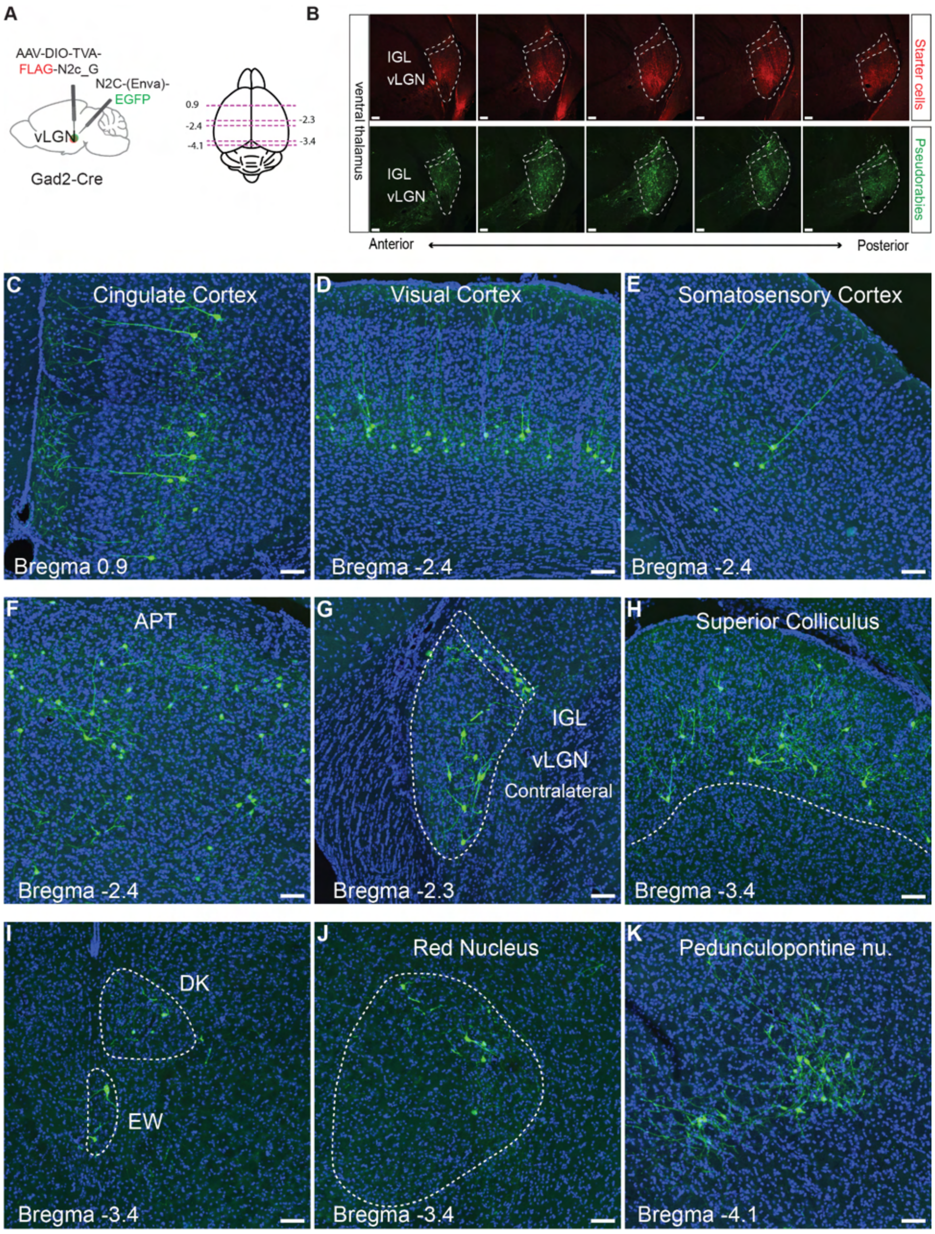


**Supplementary Fig. 1. Brain-wide projections to the vLGN reveal an integrative hub, with strong inputs from motor related regions.**

(A) Left: Brain schematic showing the location of retrograde transsynaptic vector injections in the vLGN. Right: schematic of a horizontal brain section showing the rostro-caudal locations (magenta dashed line) of the images shown in (C-K).

(B) Coronal vLGN series showing the extent of the infection. Top: starter vector expression. Bottom: pseudorabies vector expression.

(C-K) Coronal brain sections of pseudorabies transsynaptic vector expression in sensory- and motor-associated areas.

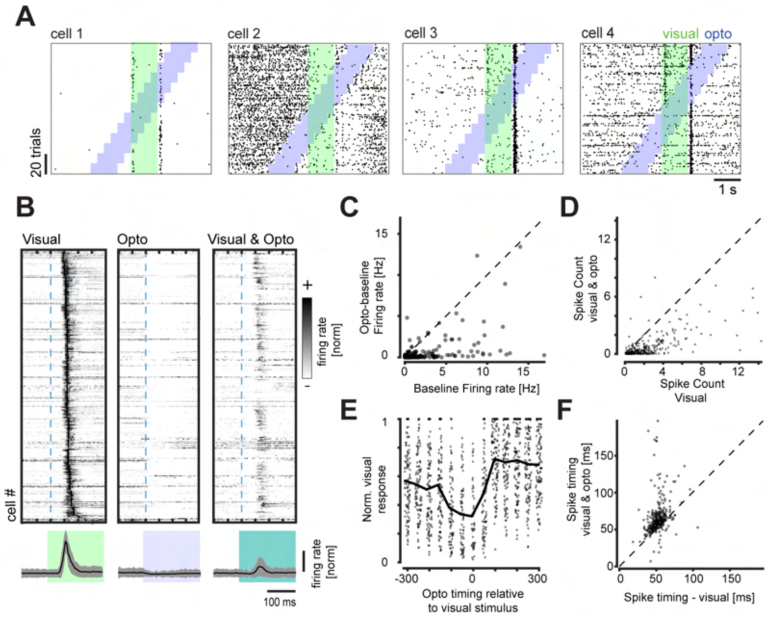


**Supplementary Fig. 2. vLGN/IGL suppression of intermediate SC *in vivo.***

(A) Spike raster plots of visually and optogenetically responsive sample neurons of the superficial SC. Visual stimulus epochs (green) are interleaved with optogenetic stimulation (blue).

(B) Sorted and normalized firing responses of superficial SC neurons to visual, optogenetic, and combined stimulation (top) with their respective mean responses (bottom, mean ± SD).

(C-D) Quantification of optogenetic baseline (C, mean ± SD of firing rate 0.09±0.21) and visually evoked suppression (D, mean ± SD of spike count 1.40±1.78) (Wilcoxon signed rank test, $p < 0.001$).

(E) Quantification of the duration of inhibition kinetics (80 units, 7 recordings, 4 animals).

(F) Analysis of the optogenetic influence on visually evoked spike timing (mean ± SD of spike timing -11.3±22.0ms, Wilcoxon signed rank test, $p < 0.001$).

390 units, 26 recordings, 5 animals if not stated otherwise

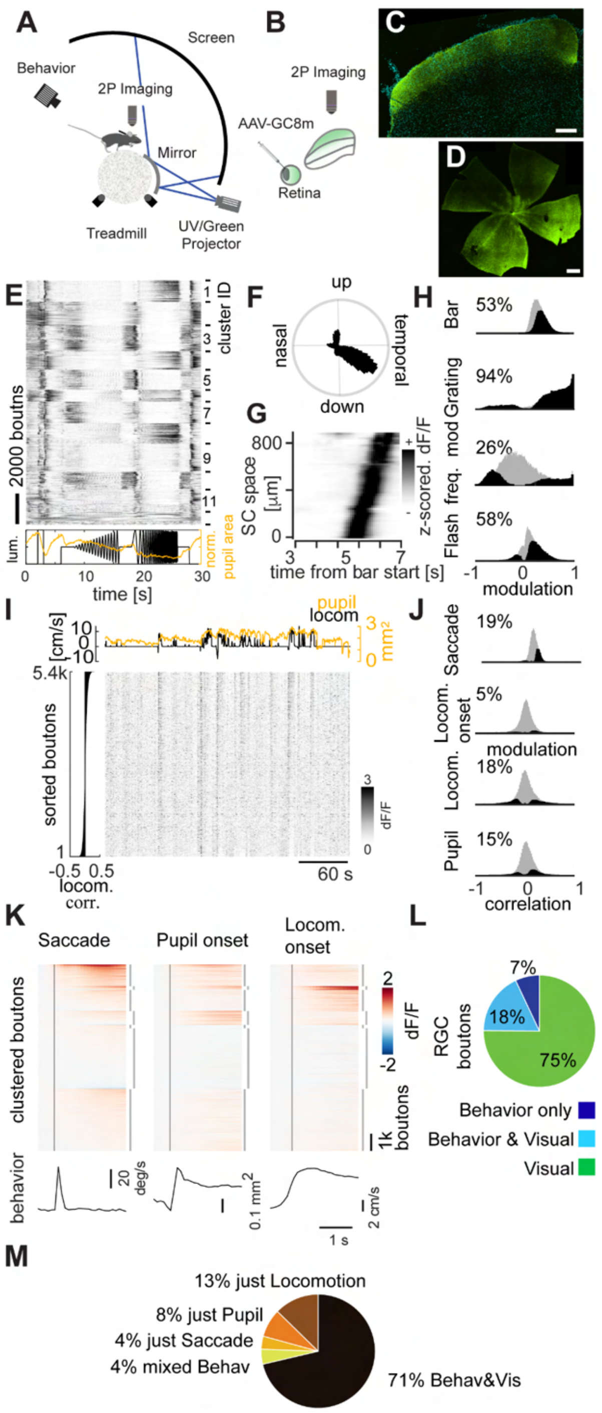


**Supplementary. Fig. 3. Retinal ganglion cell (RGC) activity in superficial SC is primarily visual**.

(A) Schematic of the 2-photon imaging setup with mice head-fixed while running on a spherical treadmill and panoramic dome visual stimulation.

(B-D) GCaMP8 was virally expressed in the retina (confocal micrograph in D, scale bar: 500 µm) and RGC axonal boutons were imaged in the sSC (confocal micrograph in C, scale bar: 200 µm).

(E) Cluster-sorted, averaged and z-scored responses (top) to full field luminance chirps (bottom, black line) of RGC boutons in the sSC and average modulation of normalized pupil area (bottom, orange).

(F) Polar histogram of direction-selective ($p < 0.01$, shuffle test, n= 40471 (64%)) RGC boutons' preferred direction in response to full field drifting gratings.

(G) Z-scored and spatially binned (20 µm) bouton responses to moving bar across SC space.

(H) Histograms of modulation indices of RGC bouton population (gray) and significantly modulated boutons (percentages, black) to different visual stimuli. Two-sided Wilcoxon signed rank test, $p < 0.01$.

(I) Raster plot of example recording of RGC boutons sorted by correlation with locomotion (left) and respective locomotion (black, top) and pupil area (orange, top) traces.

(J) Histograms of modulation indices (saccades, locomotion onset) or correlation coefficients (locomotion speed, pupil area) of RGC bouton population (gray) and significantly modulated boutons (percentages, black) to behavioral parameters. Two-sided Wilcoxon signed rank test, shuffle test for correlations (see methods), $p < 0.01$.

(K) Average activity modulation of RGC boutons (n = 40389, 19 recordings, 4 animals, only including recordings with at least 10 trials per each condition) aligned to saccades (bottom: average eye movement speed), rapid pupil expansion onset (bottom: average pupil area) and locomotion onset (bottom: average forward locomotion speed). Time of behavioral event indicated by vertical black line. Boutons sorted by k-means clustering (interrupted vertical lines on the right, k = 7) joint activity profiles.

(L) Pie chart of boutons significantly modulated by any of the visual stimuli tested (H) and/or significantly modulated by/correlated to any of the behavioral parameters (J); $p < 0.01$, Bonferroni corrected for multiple comparisons.

(M) Pie chart of boutons significantly modulated by/correlated to any of the behavioral parameters (J) individually, removing purely visually responsive neurons (n = 19866, 29 recordings, 5 animals); $p < 0.01$, Bonferroni corrected for multiple comparisons.

n = 78730, 29 recordings, 5 animals, if not stated otherwise.

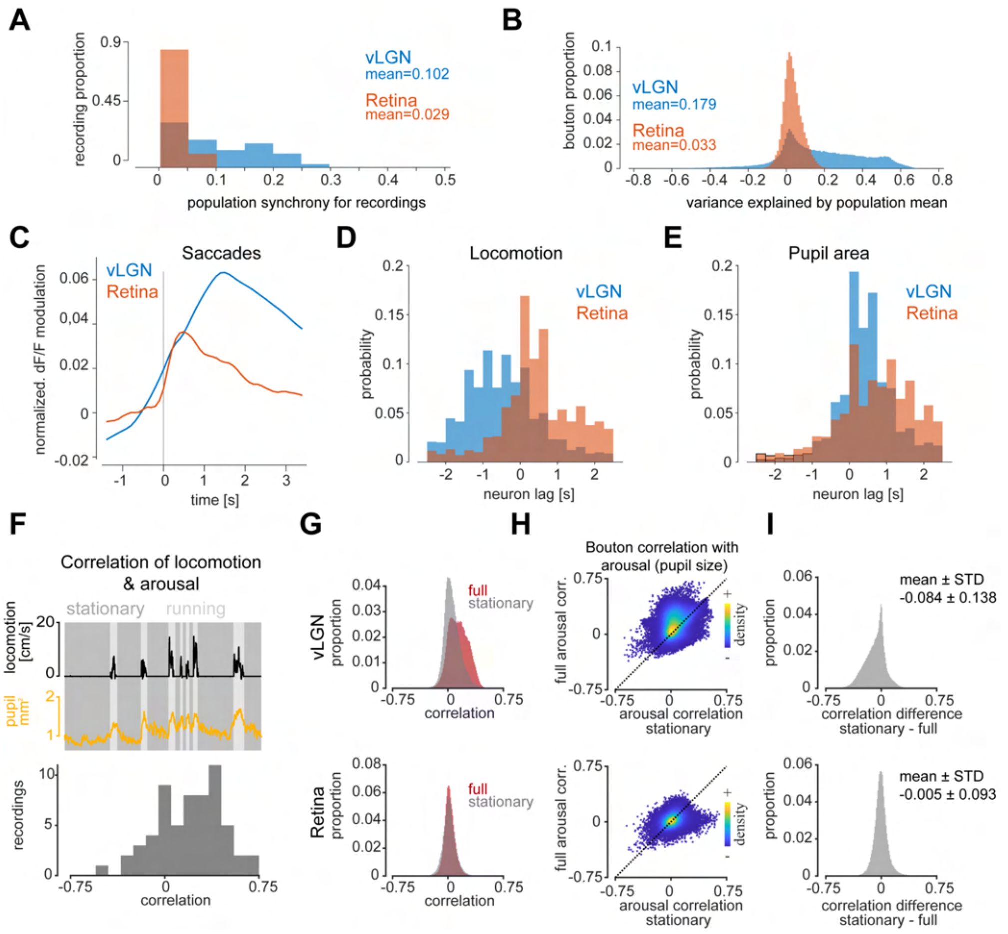


**Supplementary Fig. 4. Synchrony, timing, and behavioral dependencies of vLGN and retinal terminals in the superior colliculus.**

(A) histograms of bouton activity synchrony of vLGN/IGL (blue, n = 38 recordings) and RGC (red, n = 29) boutons within each recording.

(B) histograms of explained variance by the population mean for vLGN/IGL and RGC boutons.

(C) mean dF/F modulation of vLGN and RGC boutons following spontaneous saccades. Note: while the response kinetics are not directly comparable between the populations due to different indicators (GCaMP8m for RGCs and GCaMP6s for vLGN), the normalization provides relative contribution of saccadic responses.

(D) histograms of cross-correlation peak lags for significantly positively correlated vLGN/IGL (n = 75915, 35%) and RGC (n = 16122, 21%) boutons with locomotion speed.

(E) same as (D) but for cross-correlations with pupil area (vLGN/IGL: n = 140794, 64%; RGC: n = 15857, 20%))

(F) top, example trace of correlated locomotion (black) and pupil size (yellow). Periods of no locomotion (speed < 1 cm $s^{-1}$, “stationary”, dark gray) were selected for comparisons in (G-I). Bottom, histogram of correlation coefficients between pupil size and locomotion over pooled recordings (n = 67 recordings).

(G) normalized histograms of overall (red) correlation coefficients for vLGN (top) and RGC (bottom) boutons with pupil size as well as correlation to pupil size in stationary periods (gray).

(H) Density-colored scatter plots of bouton full correlation coefficients over stationary correlation coefficients, as in b, for vLGN (top) and RGC boutons (bottom). Two-sided Wilcoxon signed-rank test. Gray screen periods selected for (A-H).

(I) histogram of differences between stationary and full correlation coefficients, showing a strong reduction for vLGN (top) and a small reduction for RGCs (bottom).

(B-E), (G-I) vLGN: n = 220021 boutons, RGC: n = 78730.

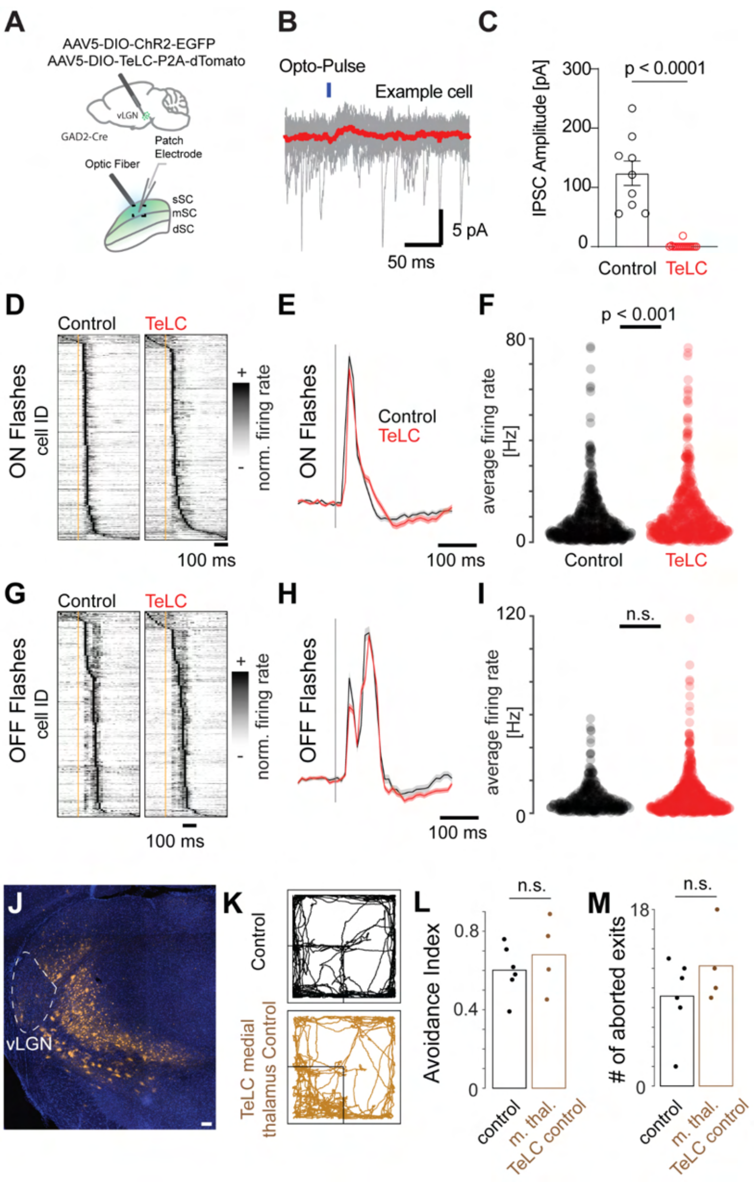


**Supplementary Fig. 5. TeLC control experiments.**

(A-C) TeLC effectively blocks synaptic transmission of the vLGN.

(A) Schematic of a combined viral infection of TeLC and ChR2 in the vLGN/IGL complex and subsequent patch clamp recordings in the sSC.

(B) Example trace of optogenetically activated IPSC in sSC neurons with TeLC block (average in red).

(C) Quantification of ChR2 evoked IPSC current amplitude with and without TeLC block (Control: 9 cells, 3 animals, TeLC: 17 cells, 2 animals; Mann-Whitney U test).

(D-I) vLGN block has a minor effect on visual responses to flashes in the SC (ON: control: 659 units, 7 recordings, 5 animals; tnt: 759 units, 10 recordings, 8 animals; OFF: control: 421 units, 7 recordings, 5 animals; tnt: 866 units, 10 recordings, 8 animals ).

(D) Normalized PSTH of mean saccade-triggered responses in control (left) and TeLC (right) mice.

(E) Normalized population-averaged saccade responses and SEM (shading).

(F) Average firing rate for the first 200 ms after saccade onset (KS test, $p < 0.001$; mean ± SD: control = 1.65 ± 1.92 , tnt = 2.01 ± 2.16).

(G-I) as (D-E) but for OFF flashes. (I) (KS test, $p = 0.2$; mean ± SD: control = 2.03 ± 2.05 , tnt = 2.34 ± 2.85)

(J-M) TeLC expression medial to the vLGN (including ZI), does not affect cliff avoidance (6 control and 4 TeLC medial thalamus control mice).

(J) Location of the viral expression for TeLC-tdTomato expression (orange) in Gad2-cre mice. Scale bar: 100 µm.

(K) Examples of running trajectories during the first 10 min for control and TeLC medial thalamus control  mice.

(L-M) Quantification of behavior per animal (Wilcoxon rank sum test).

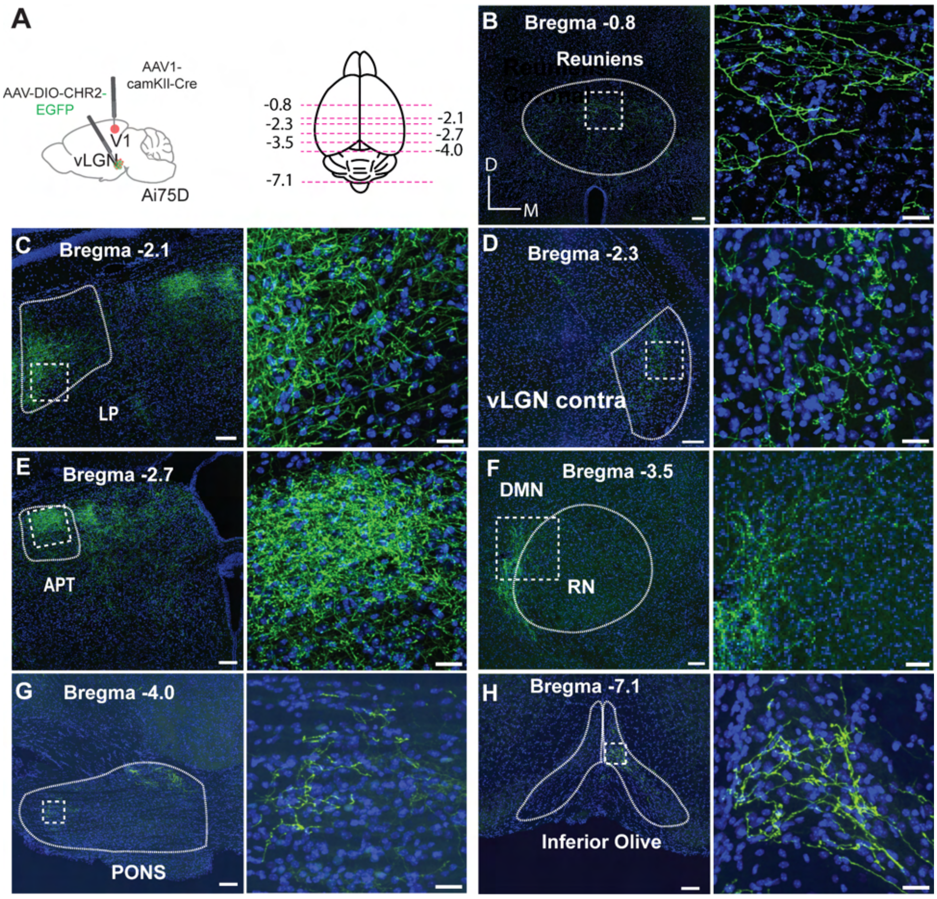


**Supplementary Fig. 6. Brain-wide vLGN projections using the AAV1-Cre intersectional approach.**

(A) Left: Brain schematic showing the location of anterograde transsynaptic vector injection. Primary visual cortex (V1, red dot) and ventral lateral geniculate nucleus (vLGN, green dot) in Ai75D mice, which express nuclear-localized tdTomato following Cre recombinase exposure. Right: Schematic of coronal confocal images of vLGN terminals.

(B-H) left: low magnification, right: close ups.

(B) Nucleus reuniens .

(C) Lateral pulvinar (LP).

(D) Contralateral vLGN.

(E) Area pretectalis (APT).

(F) Deep midbrain nucleus (DMN) and red nucleus (RN).

(G) Pons; right, high magnification of box in (E).

(H) Left, inferior olive (IO); right, high magnification of box in (F).

Low magnification images scale bars: 100 µm; close up images scale bar: 50 µm.

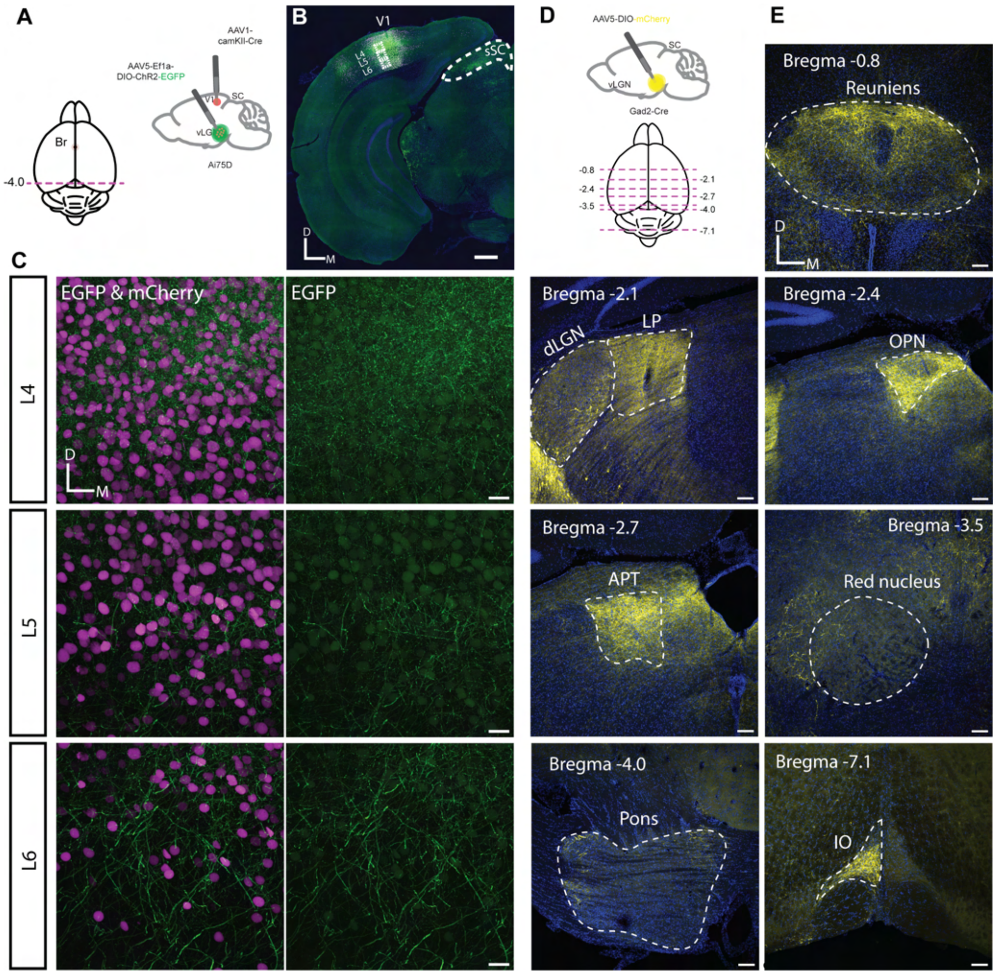


**Supplementary Fig. 7. AAV-Cre control experiments and supporting anatomical evidence of vLGN projections using Gad2-Cre expression.**

(A-C) Anterograde AAV1-Cre mediated intersectional approach, initiated in the visual cortex, is specific for the visual thalamus.

(A) Top right, a schematic of a sagittal brain showing the location of anterograde transsynaptic vector injection in the primary visual cortex (V1, red dot) and Cre-dependent vector injection in the ventral lateral geniculate nucleus (vLGN, green dot). Bottom left, a schematic of a horizontal brain section showing the rostro-caudal locations (magenta dashed line) of the images shown in (B, C).

(B) Confocal image of the injection site in primary visual cortex (V1).

(C) High-magnification confocal images of V1 across cortical layers (L4, top; L5, middle; L6, bottom). All scale bars: 20 µm, but (B) 500 µm

(D-E) Brain-wide vLGN projections using targeted infections in Gad2-Cre animals.

(D) Top right, schematic of a sagittal brain showing the location of anterograde vector injection into the vLGN; bottom left, schematic of a horizontal brain section showing the rostro-caudal locations (magenta dashed line) of the vLGN efferents and the injection site.

(E) Coronal confocal images of vLGN terminals of Nucleus reuniens, Lateral pulvinar (LP), Olivary pretectal nucleus (OPN), Area pretectalis (APT), Pons and Inferior olive (IO). Injection site in vLGN and projections to the superior colliculus are shown in Fig. 6. Scale bars: 100 µm

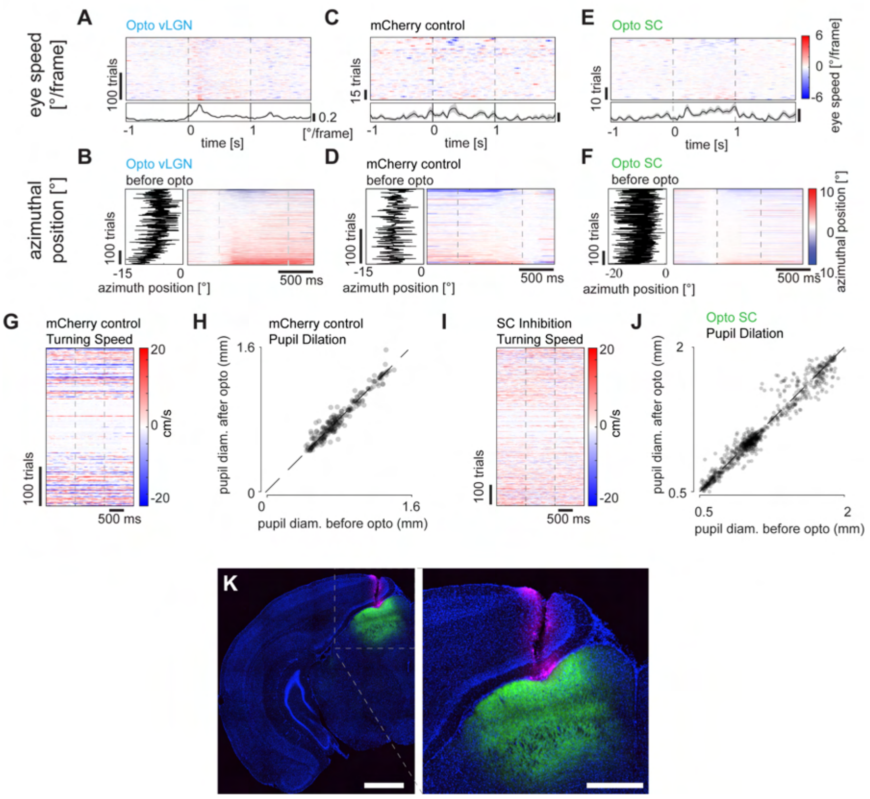


**Supplementary Fig. 8. Optogenetic behavioral controls.**

(A) Optogenetically induced saccadic eye movements sorted by mean angular speed during 0.25 s after the onset of optogenetic stimulation. Only the trials where the saccadic eye motion exceeded 14 °/s during the entire optogenetic stimulation are plotted (see also Fig. 4E). Gray dotted lines mark the start and end of the optogenetic stimulation. A clear velocity peak is visible at 150 ms post activation (211 repetitions, 9 recordings, 3 animals).

(B) Starting eye position (left) and optogenetically induced saccade (right) sorted by displacement magnitude for ChR2 experiments (544 repetitions, 11 recordings, 4 animals).

(C) As (A) but for control mCherry experiments (62 repetitions, 3 recordings, 2 animals).

(D) As (B) but for control mCherry experiments (227 repetitions, 3 recordings, 2 animals).

(E) As (A) but for SC inhibition using panneuronal eNpHR3.0 (185 repetitions, 3 recordings, 3 animals).

(F) As (B) but for SC inhibition using panneuronal eNpHR3.0 (779 repetitions, 3 recordings, 3 animals).

(G) Optogenetically induced turning speed for control mice. Gray dotted lines mark the start and end of the optogenetic stimulation (229 repetitions, 3 recordings, 2 animals).

(H) Optogenetically induced pupillary dilation for control mice (227 repetitions, 3 recordings, 2 animals, Wilcoxon signed rank test, $p = 0.88$).

(I) As (G) but for SC inhibition experiments (780 repetitions, 3 recordings, 3 animals).

(J) Optogenetically induced pupillary dilation for SC inhibition mice (779 repetitions, 3 recordings, 3 animals, Wilcoxon signed rank test, $p = 0.0002$).

(K) Optic fiber track (magenta) and eNpHR3.0 expression (green) in the SC. Scale bars: left 1mm, right 500 µm.
